# The Golgi Rim is a Precise Tetraplex of Golgin Proteins that Can Self-Assemble into Filamentous Bands

**DOI:** 10.1101/2025.03.27.645134

**Authors:** Maohan Su, Abhijith Radhakrishnan, You Yan, Yuan Tian, Hong Zheng, Ons M’Saad, Morven Graham, Jeff Coleman, Jean N. D. Goder, Xinran Liu, Yongdeng Zhang, Joerg Bewersdorf, James E. Rothman

**Author notes:** Lead contact (J.B.), (J.R.). These authors contributed equally (Maohan Su, Abhijith Radhakrishnan).

## Abstract

Golgin proteins have long been suspected to be organizers of the Golgi stack. Using three-dimensional super-resolution microscopy, we comprehensively localize the human golgin family at the rim of the Golgi apparatus at 10-20 nm resolution *in situ*. Unexpectedly, we find that the golgins are precisely organized into a tetraplex with four discrete layers, each containing a specific set of rim golgins. We observe no golgins inside the stack between its membrane-bound cisternae. Biochemically characterizing most of the golgins as isolated proteins, we find that they form anti-parallel dimers and further self-assemble into bands of multi-micron-long filaments. Based on our findings, we propose an “outside-in” physical model, the Golgin Organizer Hypothesis, in which the Golgi stack of cisternae and its overall ribbon morphology directly result from bending circumferential bands of rim golgin filaments onto a membrane surface, explaining stack formation without the need for special “stacking proteins.”

## Introduction

The defining feature of the Golgi apparatus is its stack of flattened cisternae with typically dilated rims (Klumperman, 2011). The stack is a polar structure, with compositionally distinct *cis* and *trans* faces separating by central (medial) cisternae whose composition is further distinct. In most vertebrate cells and some other metazoan cells (Benvenuto et al., 2024), these stacks join each other laterally in precise *cis*-to-*trans* register via interconnecting membrane tubules thereby forming a single topologically complex peri-nuclear continuum termed the Golgi ribbon. The numerous well-known components of vesicular transport machinery (coat proteins, SNAREs, Rab GTPases, etc.) are asymmetrically distributed across the stack thereby polarizing membrane traffic (Cai et al., 2007; Glick & Nakano, 2009; Warren & Malhotra, 1998).

As well-ordered as the structure of the stack is, it is also remarkably plastic. The ribbon is stabilized by microtubules, depolymerization of which results in reversible dissociation of the ribbon into hundreds of individual stacks, which are termed ‘ministacks’ in this instance. In cell division the ribbon naturally disassembles into ministacks, which further fragment into many thousands of smaller vesicles that efficiently distribute to the daughter cells, and then reassemble into a single ribbon within each cell (Shorter & Warren, 2002). Certain drugs that perturb vesicle traffic can trigger analogous reversible disassembly non-physiologically (Helms & Rothman, 1992; Takizawa et al., 1993). In both forms of disassembly, a distinct subset of Golgi-associated proteins termed ‘golgins’, many of which remain assembled separate from the bulk of the membrane vesicles (Puthenveedu et al., 2006; Seemann, Jokitalo, Pypaert, et al., 2000). For this reason, it has long been suspected that the golgins are organizers of the stack.

Physically, golgins are oligomeric rod-like coiled-coil proteins typically indirectly attached to the Golgi membrane bilayer at one or the other end, most often by lipid-binding adaptor proteins (myristoylated ARF or Arl or prenylated Rabs) but sometimes directly attached by a transmembrane domain. The extensive predicted coiled-coil domains of golgins are frequently interrupted, imparting segmental flexibility. Golgins are also quite long, ranging from ∼50 nm to ∼500 nm when (theoretically) fully extended (Munro, 2011); i.e., comparable in dimensions to transport vesicles and even cisternae. Their length combined with their flexibility makes them ideally suited for their well-established function as vesicle ‘tethers’, i.e., capturing vesicles for SNARE-dependent fusion with the Golgi membrane. Vesicles are captured when their surface Rab[GTP] proteins bind to a golgin tether(Barr & Short, 2003; Lowe, 2019; Witkos & Lowe, 2015). Specificity in vesicle capture results from a combination of the specific intra-Golgi localization of each golgin with its specific binding of a different Rab protein(s) and the differing Rab composition of different transport vesicles (Park et al., 2022; Sinka et al., 2008; Wong et al., 2017; Wong & Munro, 2014). The golgin family is evolutionarily conserved in eukaryotes. Vertebrates typically encode 11 canonical golgins and invertebrates like *Drosophila* or *C. elegans* encode 9 or 8 (Munro, 2011; Witkos & Lowe, 2015).

Despite their importance, little is actually known of the structure and physical chemistry of the golgins, likely because they are so large and perceived to be daunting targets for purification as full-length proteins. For example, there are only limited reports of their oligomeric state(s) and individual morphology as seen by electron microscopy (Cheung et al., 2015; Ishida et al., 2015), and these few studies have been carried out on extremely dilute (nM) solutions that do not reflect the vastly higher local concentrations at the Golgi surface (likely > 10 µM in most cases; see Discussion).

Recently, we reported that several golgins (unbound from membranes) can self-associate into condensates both as purified proteins *in vitro* (Rebane et al., 2020) and when over-expressed in cells (Ziltener et al., 2020). This work was suggested by a speculative ‘Golgin Organizer’ hypothesis (Rothman, 2019), introducing the idea that golgins could potentially phase separate from each other on the surface of the stack to self-assemble as distinct layers, each enclosing cisternae capable of capturing designated types of vesicles. In this paper we give additional clarity to the hypothesis with the finding that there may be four distinct phases at the Golgi rim, each consisting of flat bands of bundled golgin filaments.

A necessary anatomical prediction (although not proof) of this hypothesis is that the rim of each cisterna must have a unique golgin composition, representing a unique phase. Every cisterna in the stack has an exposed rim, and for the central (medial) cisternae this is their only exposed surface. It is therefore predicted (in the simplest version of the Golgin Organizer hypothesis) that each successive rim has a unique ‘Rim Golgin’ or unique combination of golgins. The *cis* and *trans*-most cisternae of course will also have additional surfaces exposed, and these will have yet additional non-rim golgins. which are termed ‘Sheet Golgins’. A measurable prediction is the sharp boundaries between golgins.

Unfortunately, suitable quantitative compositional measurements with single cisternal resolution have been out of reach due to technical reasons. Certainly, electron microscopy (EM) has single cisternal resolution, but molecular detection involves variable and unknown undercounting due to sparse antibody sampling, even when suitable antibodies are available. Optical detection is both more versatile and quantitative, but diffraction-limited microscopy lacks the resolution to resolve individual cisternae. Even conventional (one objective) super-resolution methods fall short because while they can have suitable spatial resolution (∼ 25 nm) in the plane of focus (X-Y), they have inadequate resolution for the Golgi stack in depth (Z), typically in the ∼70-100 nm range, greater than the center-to-center distance between neighboring cisternae. Those limitations give rise to the appearance of a smooth gradient distribution of Golgins and other Golgi-resident proteins along the cis-trans axis, even if they were discretely distributed to single cisternae. Important progress has been made by taking advantage of the favorable geometry of side (X-Y plane) views of nocodazole-induced ministacks (Tie et al., 2018, 2022) combined with extensive averaging to offset variations from ideal views. But even here the single cisternal resolution needed to rigorously test the Golgin Organizer hypothesis has been unachievable.

A step-change advance in super-resolution microscopy was whole-cell 4Pi single-molecule switching (4Pi-SMS) microscopy, which uses a second objective to enable precise interferometric determination of a molecule’s Z-position, thereby affording practical isotropic resolution in fixed whole cell samples of 10-20 nm (in X, Y, Z) (Huang et al., 2016), more than sufficient for single-cisterna resolution in individual Golgi ribbons and ministacks, without averaging. The recent addition of multi-color imaging by salvaged fluorescence eliminates chromatic shifts and enables precise quantitative colocalization analysis (Y. Zhang et al., 2020).

Here, we set out to use those new methods to investigate the intrinsic physical properties of human golgins, and to establish their precise localization within the three-dimensional (3D) structure of the entire native Golgi ribbon in human cell lines. In this work, in addition to tools of protein biochemistry, we used 4Pi-SMS super-resolution microscopy to systematically image golgins and for reference and validation some other Golgi-resident proteins as well. These results were cross-checked with pan-staining expansion microscopy (Pan-ExM) and immuno-electron microscopy (immuno-EM) where possible. We find that the rim of each cisterna has a unique golgin composition, sometimes just a single rim golgin and in other cases a unique combination of several rim golgins. Remarkably, the rim golgins share a common intrinsic biochemical property by self-assembling into multi-micron long filaments similar in dimension to the perimeter of the intact Golgi apparatus.

## Results

### Individual golgins are located either at cisternal rims or sheets of the Golgi

In this study our aim was to systematically localize most endogenously expressed golgins using specific antibodies and a secondary antibody for detection. To achieve single-cisterna precision, we employed 4Pi-SMS microscopy with salvaged fluorescence. We focused on the perinuclear Golgi ribbon in paraformaldehyde-fixed interphase HeLa cells, HUVEC and SAOS2 cells. We localized a total of 10 different golgins, typically in combinations with other golgins for direct comparison. Table S1 lists the golgins we have localized, and the source/properties of the antibodies used.

The golgins clearly fell into two categories, as expected from earlier reports using imaging approaches affording lower resolution (Harada et al., 2024; Tie et al., 2018; Witkos & Lowe, 2015). The majority were located only at the periphery of the ribbon, delineating the rims of the structure. These are termed ‘rim golgins’. The others were located within the ribbon itself in regions that correspond to the extensive sheet-like flat faces of cisternae. These are termed ‘sheet golgins’ and will be shown later to be located at either the *cis* face or *trans* face of the Golgi stack.

Fig. 1A (Movie S1) presents an example of two-color 4Pi-SMS images showing that the rim golgin Giantin (*GOLGB1*) decorates the rim of a sheet, while the sheet golgin GM130 (*GOLGA2*) lies entirely within a convoluted but essentially flat surface that can be traced through 3D space. The isotropic resolution afforded by 4Pi-SMS enables us to freely rotate the point cloud data without elongation artifacts to obtain the complete 3D image. Though such 3D data are available in all cases, for simplicity we generally showed only two approximately orthogonal zoomed-in views: *en face* (looking down on the top of sheets) and a *side view* analogous to a cross-section in conventional EM (Fig. 1C). Line profiles across the span of an *en face* view of a sheet confirms complete separation of Giantin (rim) from GM130 (sheet), suggesting a clear segregation without a concentration gradient (Fig. 1B, left). The line profile across a side view showed a shift in peak intensity, indicating that Giantin is not in the same cisternal sheet as GM130 (Fig. 1B, right). As will be seen later, they are in fact in successive cisternae. The rim localization of Giantin at the Golgi apparatus from our data is consistent with previous immuno-EM and Airyscan confocal microscopy (Koreishi et al., 2013; Tie et al., 2018).

**Figure 1.**
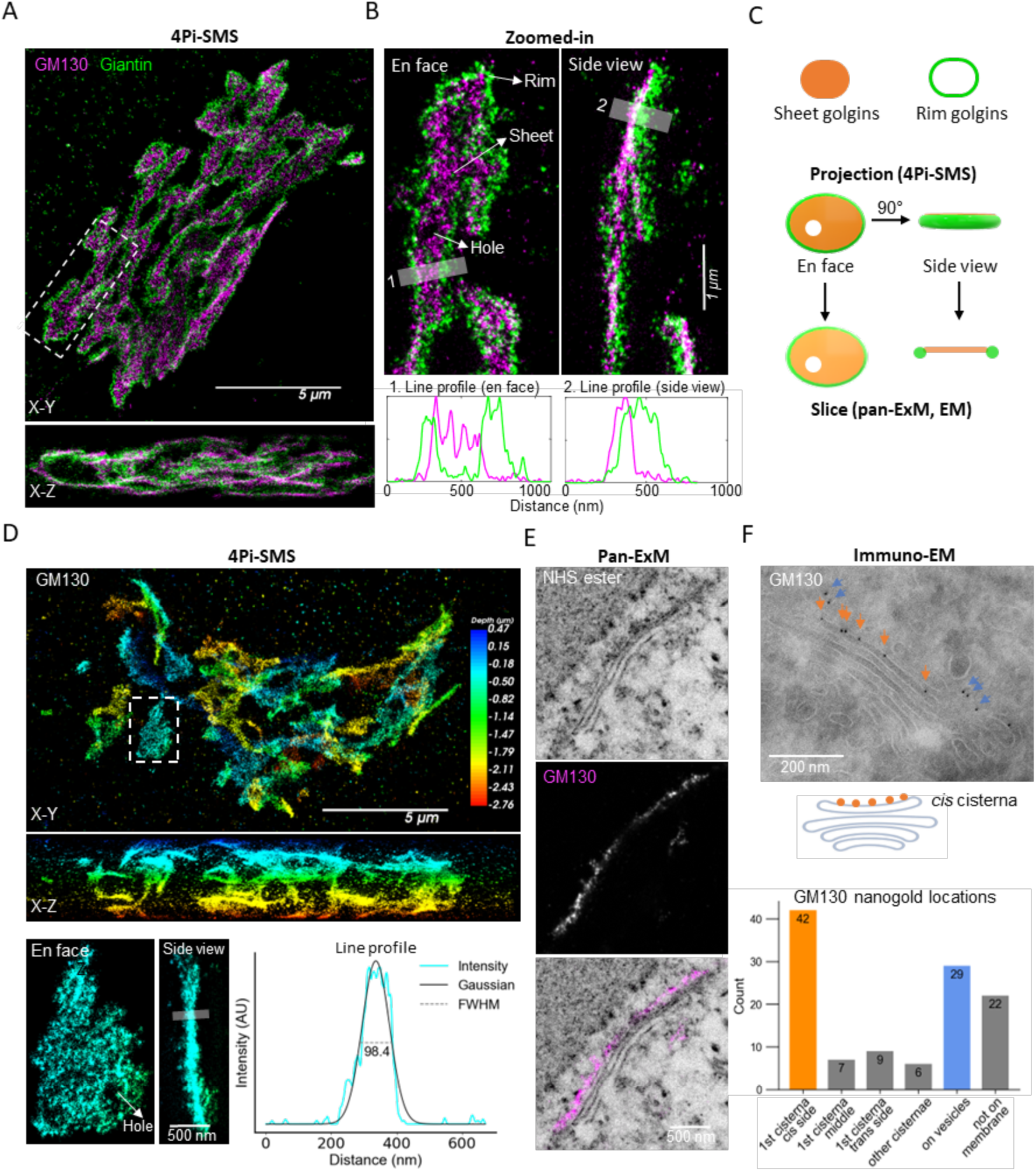
Superresolution 4Pi-SMS microscopy exhibits segregation of sheet and rim golgins. A. Overview of 4Pi-SMS two-color images of immuno-labeled endogenous GM130 (BD, mouse) and Giantin (Sigma, rabbit) in HeLa cells shown in X-Y (Top) and X-Z view (Bottom). Also see Movie S1. B. Zoom-in region (outlined by white dashed box in A) shown in both *en face* (Left) and side view (Right) with Line-scan profiles (Bottom). C. Illustration of presentations of sheet and rim golgins used in this work. D. 4Pi-SMS image of GM130 in HeLa cells as shown in X-Y view (Top) and X-Z view (Middle). Color denotes depth. Bottom: Zoomed-in images of the white dashed box (Left) with line-scan plot and Gaussian fitting (Right). E. Pan-ExM images of NHS ester (Top), GM130 (Middle) and merge (Bottom). Scale bars are corrected for the expansion factor. F. Top: Electron microscopy image of GM130 immuno-labeled with 10 nm nanogolds. Middle: Illustration of GM130 (orange dots) on the cis side of cis cisterna. Bottom: Distribution of 114 nanogold particles in 13 Golgi Apparatus.

The sheet golgin GM130 is present in distinct 30-60 nm diameter puncta which are located on a common and continuous sheet-like convoluted surface (Fig. 1D). GM130 is known from previous work to be indirectly bound to the membrane recruited by N-myristoylated GRASP proteins (Barr et al., 1998; Yoshimura et al., 2001). As expected, GM130 and GRASP65 are co-localized within the puncta (Fig. S1A).

The sheet is occasionally perforated with openings of 50-500 nm in diameter within the otherwise continuous surface (Fig. 1B, D, 2A, S1A arrows). These openings resemble in size, frequency and shape of the fenestrations that had previously only been revealed through electron tomography (Ladinsky et al., 1999). Some large openings are aligned in *cis*-*trans* direction, penetrating through multiple stacked cisternae, and thus are bounded within the stack by internal cisternal rims (for example Fig. 3B arrows). It is notable that the internal rims of the fenestrations do not have Giantin (or as will be seen any other rim golgin) (Fig. 1B, 2A); rim golgins are only found at the true external rim that defines the outer contour of the Golgi ribbon as a whole, indicating mechanisms beyond simple curvature preferences.

**Figure 2.**
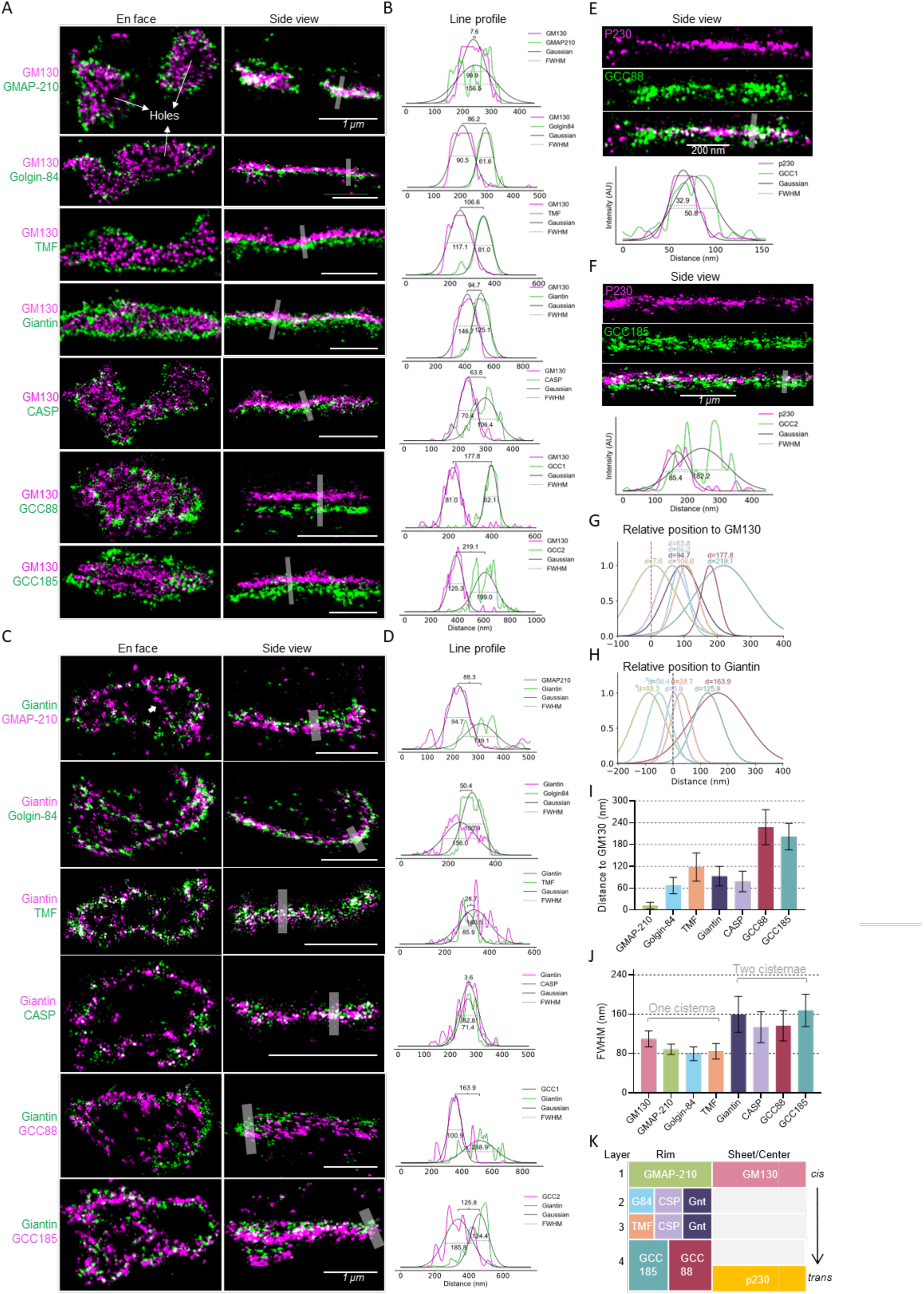
Pairwise imaging reveals tetraplex layer organization of rim golgins. A. Representative 4Pi-SMS images of zoomed-in Golgi stacks labeled with GM130 and different rim golgins. Both *en face* (Left) and side view (Right) are shown. Also see Fig. S2, S3, Movie S2, S3. B. Corresponding line-scan plots of the lines drew in A. Plots were Gaussian fitted for each channel, and FWHM and the distance between the peaks were written on graphs. C. Representative 4Pi-SMS images of zoomed-in Golgi stacks labeled with Giantin and different rim golgins. D. Corresponding line-scan plots of the lines drew on the left images in C. Plots were Gaussian fitted for each channel, and FWHM and the distance between the peaks were written on graphs. E. Side view of GCC88 with p230 and line-scan of the line drew on the image. F. Side view of GCC185 with p230 and line-scan of the line drew on the image. G. Aligned plots of all Gaussian-fitted curves in B. Dashed line represents center positions of GM130. Colors represent each golgin as in K. H. Aligned plots of all Gaussian-fitted curves in D. Dashed line represents center positions of Giantin. Colors represent each golgin as in K. * denotes added negative sign based on information from I and J to show GMAP-210 and Golgin-84 are on different side of Giantin. I. Statistics of peak-to-peak distance from various rim Golgins to GM130. N=13-33 cisternae from 5-11 cells for each protein. See also Table S2. J. Statistics of FWHM from various Golgins. N=10-16 cisternae from 4-8 cells for each protein. See also Table S2. K. Diagram of relative positions of Golgins on the cross-section of a Golgi stack inferred from all the data above.

**Figure 3.**
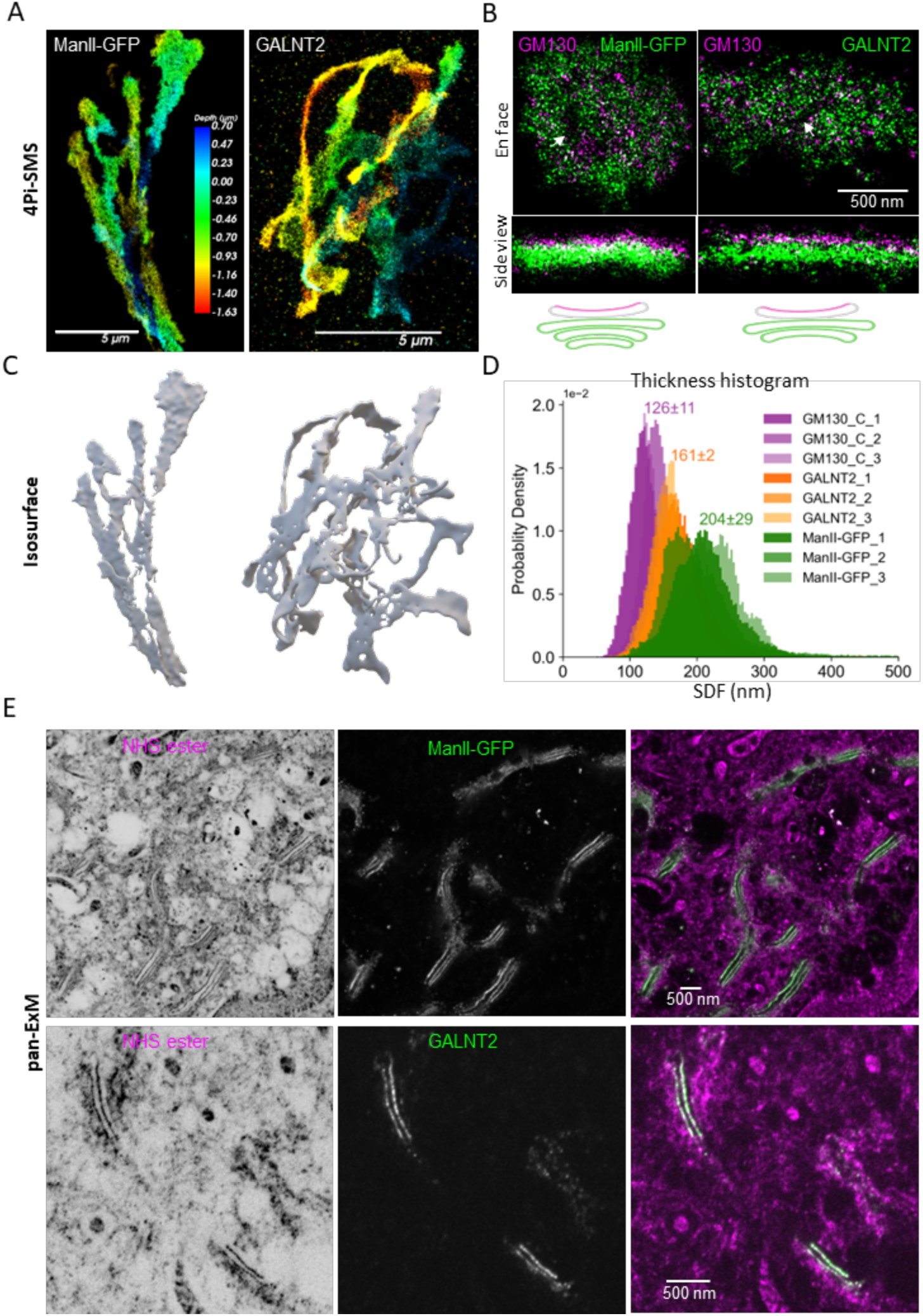
ManII-GFP and GALNT2 occupy multiple sheets of medial cisternae. A. 4Pi-SMS images of overexpressed ManII-GFP (Left) and GALNT2 (Right) in HeLa cells. Color denotes depth. B. Two-color images of ManII-GFP (Left) and GALNT2 (Right) with GM130. Arrows show aligned holes in multiple cisternae. Bottom illustrates our interpretation of different thickness between ManII-GFP and GALNT2 in a side view. C. Shrink-wrapped isosurface of ManII-GFP (Left) and GLANT2 (Right). See Fig. S5. D: Histogram of isosurface thickness of GM130, GALNT2, ManII-GFP (three cells each). SDF is shown on top as mean±SD. E. Top panel: Pan-ExM images of NHS ester (Left), overexpressed ManII-GFP (Middle) and merge (Right). Bottom panel: Pan-ExM images of NHS ester (Left), GALNT2 (Middle) and merge (Right).

### The sheet golgin GM130 is located exclusively on the exposed *cis* face of the Golgi stack

GM130 has been previously located by immuno-EM to the *cis*-most (first identifiable) cisterna of the stack (Nakamura et al., 1995). However, the low density of labelling by gold particles precluded determining whether GM130 was located on the cis-most (cytoplasm-facing) free surface of the first cisterna, or instead on its internal (stack-facing) surface, or possibly on both surfaces. Using 4Pi-SMS together with pan-ExM and immuno-EM has enabled us to make this important distinction.

We took line profiles across side views of GM130 sheets and calculated the (full) width of their intensity distributions at half-maximum (FWHM) from Gaussian-fitted curves. The overall measured thickness of GM130 was 110±16 nm (33 ribbons from 11 cells; Fig. 1D). This value exceeds substantially the resolution of the used 4Pi-SMS microscope; a combination of the Golgi ribbon being slightly curved across the depth of the 3D data set over which the side view is projected, the range of protein orientations, and the antibody size, all are expected to cause a broadening of the profile. The measured thickness is therefore consistent with GM130 being localized at one or two cisternae.

To narrow this further down, we used Pan-ExM to study the localization of GM130 in the context of whole Golgi stacks. This technique isotropically expands fixed cells ∼15-20 fold with a permeating expanding hydrogel while preserving most antigens. The proteome is stained by reaction of densely available NH_2_ groups with an N-hydroxy-succinimide (NHS)-dye conjugate, allowing 3D views of well-resolved individual Golgi cisternae and other ultra-structures reminiscent of conventional EM sections from confocal microscopy (M’Saad & Bewersdorf, 2020). Pan-ExM cross-sections confirm that Giantin is located at the rim of Golgi cisternae (Fig. S1C) and that GM130 is present in the sheet region (Fig. 1E). Compared with the NHS ester channel which reveals multiple cisternae, GM130 is located only in the first cisterna (Fig. 1E). Many such images suggest that it resides only at the exposed (non-stacked) surface. To further confirm this observation, we employed immuno-EM. The high density of labelling in our study fortunately enabled quantitative analysis which confirmed that the nanogold particles were much more likely to be located on the exposed *cis* face of the first cisterna facing the cytosol rather than in-between the stacks (Fig. 1F orange arrow, and Fig. S1B).

Combined, our observations strongly suggest that GM130 is located at the *cis*-side of the first (*cis*) cisterna and not in between cisterna 1 and 2. This molecular configuration aligns with the functions of its N-terminus in vesicle tethering through p115 (Nakamura et al., 1997; Seemann, Jokitalo, & Warren, 2000), and linking with microtubule nucleation through AKAP450 (Rivero et al., 2009), but does not appear to support a recent model that requires GM130 to be at rims to link Golgi stacks laterally (Y. Zhang & Seemann, 2024).

### The rim golgins are located in four discrete layers along the *cis*-*trans* axis of the Golgi

To determine the location of rim golgins along the *cis*-*trans* axis we examined 4Pi-SMS side views of Golgi ribbons stained for GM130 to locate the *cis* face and stained for the rim golgin of interest with a second color. Fig. 2A shows representative zoomed-in single ribbon views for each rim golgin. From the Gaussian-fitted intensity profiles of side views we found that rim golgins had differing peak-to-peak separations from GM130 (Fig. 2B, G). GMAP-210 (*TRIP11*) colocalized with GM130 (Movie S2) in what we term ‘Layer 1’. Giantin (*GOLGB1*), CASP (*CUX1*), Golgin-84 (*GOLGA5*), TMF (*TMF1*) were axially separated by ∼100 nm from GM130. GCC88 (*GCC1*) and GCC185 (*GCC2*) (Movie S3) were separated by ∼200 nm. This segregation implied that rim golgins localize in three main successive groups: *cis* (GMAP-210), *medial* (Giantin, CASP, Golgin-84, TMF) and *trans* (GCC88, GCC185). The segregation was confirmed by a similar imaging strategy using Giantin as the reference (Fig. 2C). The line profile showed ∼100 nm shift of GMAP-210, overlap of CASP, partial overlap for Golgin-84 and TMF, and >100 nm shift from GCC88 and GCC185 (Fig. 2DH), confirming the working model of *cis*, medial, *trans* separation we made with GM130 as the reference.

GMAP-210 had been shown to be a *cis* golgin through colocalization with GM130 in confocal microscopy data (Cardenas et al., 2009; Infante et al., 1999) and upon overexpression in immuno-EM (Pernet-Gallay et al., 2002) but the rim localization has not been reported previously, except for the *Drosophila* homolog (Friggi-Grelin et al., 2006). To confirm, we used another antibody against different regions of GMAP-210 and obtained similar results (Fig. S2A). It is noteworthy that, in addition to its rim localization, GMAP-210 sometimes had a line-like distribution across a sheet region, without aligning with other rim golgins (Fig. 2C, S2B, C, arrows). This may indicate segregated rod-like regions on or within the *cis* face’s membrane.

We observed that the medial rim golgins exhibited different distributions along the *cis-trans* axis. Giantin and CASP showed a broader *cis*-*trans* distribution (FWHM) than either TMF or Golgin-84 and Giantin fully includes both (Fig. 2D, H). Since TMF is further separated along the *cis*-*trans* axis from GM130 than Golgin-84 is (Fig. 2B, G), we concluded that the Giantin-containing grouping is sub-divided into two distinct layers, one containing Golgin-84 (‘Layer 2’) and the other containing TMF (‘Layer 3’). Previous immuno-EM studies did not obtain a precise localization for Golgin-84 (Satoh et al., 2003) or a specific cisterna for TMF (Yamane et al., 2007).

The two *trans-most* rim golgins GCC88 and GCC185 localize similarly along the *cis*-*trans* axis relative to GM130 (Fig. 2A, B, Fig. S4A, B) or Giantin (Fig. 2C, D) in ‘Layer 4’. To make finer distinctions we used the *trans*-face sheet golgin p230 (thin region, see below) as a closer point of reference. GCC88 and GCC185 could not be resolved from each other, but the layer they both occupy (Layer 4) is thicker and almost double the width of the distribution of p230 (Fig. 2E, F). To further explore if any difference in the localization of GCC88 and GCC185 could be identified, we localized them simultaneously in the same cell. Due to species limitations on primary antibodies, in this case we expressed GCC88 containing a FLAG tag and selected low-expressing cells to image it together with endogenous GCC185. The two golgins precisely co-localized in a side view (Fig. S4C) confirming their joint presence in Layer 4. However, *en face* views revealed that the epitopes detected in GCC88 and GCC185 were located in distinct radial sub-regions emanating from the outer rim of Layer 4, GCC88 being closer to the center and GCC185 further removed. Previously these two golgins had been shown to localize to distinct sub-regions of the *trans* Golgi by immuno-EM (Derby et al., 2004; Luke et al., 2003).

In summary (Fig. 2K), we observe that the golgins at the rims of the Golgi stack are organized into four distinct layers (‘Golgin Layers’). Layer 1 consists of GMAP-210. Layer 2 consists of a mixture of Golgin-84, CASP and Giantin. Layer 3 consists of a mixture of TMF1, CASP and Giantin. Layer 4 consists of a mixture of GCC88 and GCC185.

### Each rim Golgin Layer typically surrounds a single membrane cisterna except at the *trans* face

The traditional tool of viewing compartmental organization in the Golgi stack is EM, which reliably visualizes the cisternal membranes, but does not show the golgins at their periphery. From this perspective, Golgi stacks contain variable numbers of cisternae depending on cell type. In most animal cells there is a minimum of 4 cisternae, but as many as 11 has been reported (Klumperman, 2011; Rambourg & Clermont, 1997). It is important to correlate our seemingly invariant tetraplex (four) layer organization of rim golgins to the more variable number of cisternae observed by membrane visualization. The simplest possibility is that some Golgin Layers contain more than one cisterna. For example, when there are 4 cisternae, each layer would enclose a single cisterna. But when there are 5 cisternae, one or another Golgin Layer would contain 2 membrane cisternae, and so on.

As one way to test this hypothesis, and then to determine which Golgin Layers can contain variable numbers of cisternae, we compared the average thickness (FWHM) of each layer and the average center-to-center separations between them when possible (Z_Golgins_) with the average center-to-center spacing between the cisternae in the stack Z_C-C_ = 64±16 nm (M’Saad & Bewersdorf, 2020). In this idealized representation, Z is measured as the peak-to-peak distance between the two Gaussian-shaped curves fitted to numerous individual line profiles taken along the *cis-trans* axis of pairwise Golgin Layers. Our results show discretely localized rim golgins (Table S3). The thickness (FWHM measurements) of GMAP-210 (89±11 nm), Golgin-84 (80±14 nm), TMF (85±16 nm) is consistent with a single cisterna (Fig. 2J). In comparison with Z_C-C_ = 64±16 nm, the observed values for Z_GM130-GMAP210_ (11±9 nm), Z_GM130-G84_ (65±20 nm), Z_GM130-TMF_ (126±30 nm) are compatible with being sequentially separated by one cisterna (Fig. 2I). On the other hand, the thickness of Giantin (159±37 nm), CASP (134±32 nm) GCC88 (137±31 nm), GCC185 (168±33 nm) is about twice as large, and is more variable, suggesting these golgins cover the rims of two cisternae (Fig. 2J). Z_GM130-Giantin_ (93±27 nm) and Z_GM130-CASP_ (78±28 nm) are in between the values for Z_GM130-G84_ and Z_GM130-TMF_, consistent with Giantin and CASP occupying both the cisternae of Golgin-84 and TMF. The large values of Z_GM130-GCC1_ (228±48 nm) and Z_GM130-GCC2_ (202±36 nm) indicate two more cisternae after TMF.

Altogether, this quantitative analysis suggests a simple model in which each of Layer 1-3 covers a single membrane cisterna, and Layer 4 covers two or more cisternae (Fig. 2K). Larger Golgi stacks (>5 membrane cisternae) would thus contain a further expanded Layer 4 at the *trans*-face. It appears that the rim Golgin Layers are a stereotyped feature of Golgi compartmental organization, even more so than the number of cisternae they enclose.

### GALNT2 and ManII-GFP occupy exactly 2 and 3 medial membrane cisternae

While we readily found golgins localized as extended two dimensional sheets at the exposed *cis* and *trans* faces of the Golgi, none of the many golgins we studied were found internally. To rule out that this was somehow the artefactual result of a lack of antibody penetration into the center of Golgi stack, we GFP-labeled oligosaccharide processing enzyme ManII (alpha-mannosidase 2) and antibody-labeled GALNT2 (Polypeptide N-acetylgalactosaminyltransferase 2), which were reported to be primarily localized within several *medial* cisternae (Cosson et al., 2002; Novikoff et al., 1983; Rabouille et al., 1995; Röttger et al., 1998). ManII and GALNT2 are both globular membrane proteins and the epitopes of GALNT2 targeted by the antibodies we used are located within the lumen of cisternae. If antibodies can access such epitopes within the lumen, they would certainly be expected to access golgins in the cytoplasmic spaces between cisternae, were they present there.

As expected, overexpressed ManII-GFP and endogenous GALNT2 both show continuous, sheet-like localization (Fig. 3A). Both colocalize with GM130 *en face* but appear shifted in the side view (Fig. 3B), indicating their location in adjacent cisternae. However, we were not able to resolve separated multiple cisternae of ManII-GFP or GALNT2 but instead observed one thick stack. This is similar to Giantin for which we rarely were able to resolve two individual rim layers. We reason that it is possibly due to imperfect fixation at nanometer scale by paraformaldehyde (Tanaka et al., 2010).

Having ruled out the potential artifact with respect to the lack of internal golgins, we sought to use the new methods to better assign specific cisternae to each processing enzyme than was possible in previous reports. For this purpose we used the “shrink-wrapping” approach to generate accurate 3D surfaces from point-cloud data (Marin et al., 2023) (Fig. 3C, Movie S3) and estimated their “thickness” by calculating the shape diameter function (SDF) of every vertex on the isosurface (see Methods and Fig. S5; Shapira et al., 2008). We first measured GM130 as a reference and the histogram of its SDF peaked at 126±11 nm, similar to our previous manual FWHM measurement. The isosurface thickness of GALNT2 and ManII-GFP was measured as 161±2 and 204±29 nm, respectively (Fig. 3D). The ∼40 nm difference between them is consistent with one additional cisterna.

Considering that GM130 occupies one (*cis*) cisterna (Fig. 1D-F), we inferred that GALNT2 occupies two (2^nd^ and 3^rd^) and ManII-GFP occupies three cisternae (2^nd^ to 4^th^). To confirm this, an imaging method that can discern individual cisternae is needed. We chose pan-ExM and reasoned that since the protein density inside the cisternal lumen was higher than in the intracisternal space (Engel et al., 2015), the hydrogel expansion might be greater in between cisternae, exaggerating cisterna separation in post-expansion samples. Indeed, images from pan-ExM of GALNT2 and ManII-GFP showed exactly 2 and 3 layers, respectively, in most Golgi stacks checked (Fig. 3E, Movie S4).The number of layers of ManII-GFP was consistent with our previous pan-ExM data (M’Saad & Bewersdorf, 2020). The total cisterna number in the NHS-ester channel varied from 4 to 7, indicating that the cisterna number variation between different Golgi stacks most likely arises from *trans* Golgi/TGN, which remains to be tested. The pan-ExM data support the findings from the isosurface thickness measurements and exemplify the differential distribution patterns of glycol-processing enzymes within the Golgi apparatus. The unequivocally confined localization here stands in contrast to previous immuno-EM studies which showed a much wider Gaussian-like distribution of ManII or GALNT2 from *cis* to *trans* (Cosson et al., 2002; Novikoff et al., 1983; Röttger et al., 1998), probably because of the difference in labeling efficiency.

### Cross-validation of assignments of rim golgins to specific medial cisterna

An independent registration can be achieved by visualizing the rim Golgin Layers simultaneously with GALNT2, now that we know this membrane enzyme is found in precisely two medial cisternae. *En face*, Giantin is located at the rims of the GALNT2-containing cisternae which is found throughout the flat cisternal sheets (with some holes and fenestrations) (Fig. S6A, top). From the side view, Giantin and GALNT2 have similar FWHM values with a very small (∼15 nm) peak-to-peak distance, as both are expected to be on the 2^nd^ and 3^rd^ cisternae in the stack (Fig. S6A, bottom). In contrast, TMF also occupies the rim of GALNT2 sheets, but it only partially colocalizes in a side view and has about half the FWHM of GALNT2, consistent with occupying only the 3^rd^ cisterna in the stack (Fig. S6B).

Using a similar argument, TMF (Layer 3) is predicted to be closer to the *trans* Golgi than Golgin-84 (Layer 2). Indeed, using p230 as the reference, 4Pi-SMS showed that both TMF and Golgin-84 encircle p230 *en face* but appear shifted in a side view. As expected, there is an obvious gap between Golgin-84 and p230 (white arrows) (Fig. S6D, bottom) but not for TMF (Fig. S6C). These data independently confirm the medial*-trans* organization of the Golgi and establish the validity of our mapping results as illustrated in Fig. S6E.

### Golgin-97 and Golgin-245/p230 localize to the *trans* face

The *trans* face of the Golgi stack, often termed the *trans*-Golgi network (TGN) consists morphologically of one or more highly convoluted cisternae from which a network of tubules emanate. It is from here that vesicles and tubules targeted to multiple destinations emerge, and where recycling vesicles and tubules return to (Bonifacino & Rojas, 2006; De Matteis & Luini, 2008; Di Martino et al., 2019). Golgin-245/*trans*-Golgi protein p230 (*GOLGA4*) has previously been localized to *trans* cisternae using immuno-EM (Kooy et al., 1992). Golgin-97 (*GOLGA1*) shares with p230 a GRIP domain by which both are recruited to *trans* cisternae by binding to Arl1[GTP] (Lu & Hong, 2003; Munro & Nichols, 1999). Concordantly, they both localize to the sheet-like domains at the *trans* face, as we have shown here using 4Pi-SMS. In the side view, Golgin-97 is separated from GM130 by a clear gap, which presumably is due to a lack of labelling of the two medial cisternae (Fig. 4A). P230 shows a similar pattern (Fig. 4B). Golgin-97 also colocalizing with the membrane protein TGN46 (Fig. S7B). Both *trans* golgins appear to have thicker profiles in side-views than GM130 in certain regions. On the other hand, our previously published measurements of line profiles of manually selected regions using the same 4Pi-SMS microscope showed a similar thickness between p230 and GM130 (Y. Zhang et al., 2020). To resolve this potential discrepancy and objectively quantify the thickness of Golgin Layers in all regions of the Golgi apparatus in a statistically meaningful way, we again used the isosurface approach and measured the isosurface thickness by SDF. We noticed that p230 and Golgin-97 form complementary nano-domains as seen in *en face* views (Fig. 4C, S7A), consistent with previous work (Derby et al., 2004). Since neither Golgin-97 nor p230 marked the whole *trans* face, we imaged both proteins in the same cell and combined this data for the isosurface generation. The SDF histogram of p230 + Golgin-97 has a peak value of 136±4 nm similar to GM130, but with a much longer tail on the right, accounting for roughly 1/3 of data (Fig. 4D, arrow). This observation suggests that at least some portion of *trans* sheet cisternae is thicker than one layer, whereas other regions consist of a single cisterna (corresponding to the peak).

**Figure 4.**
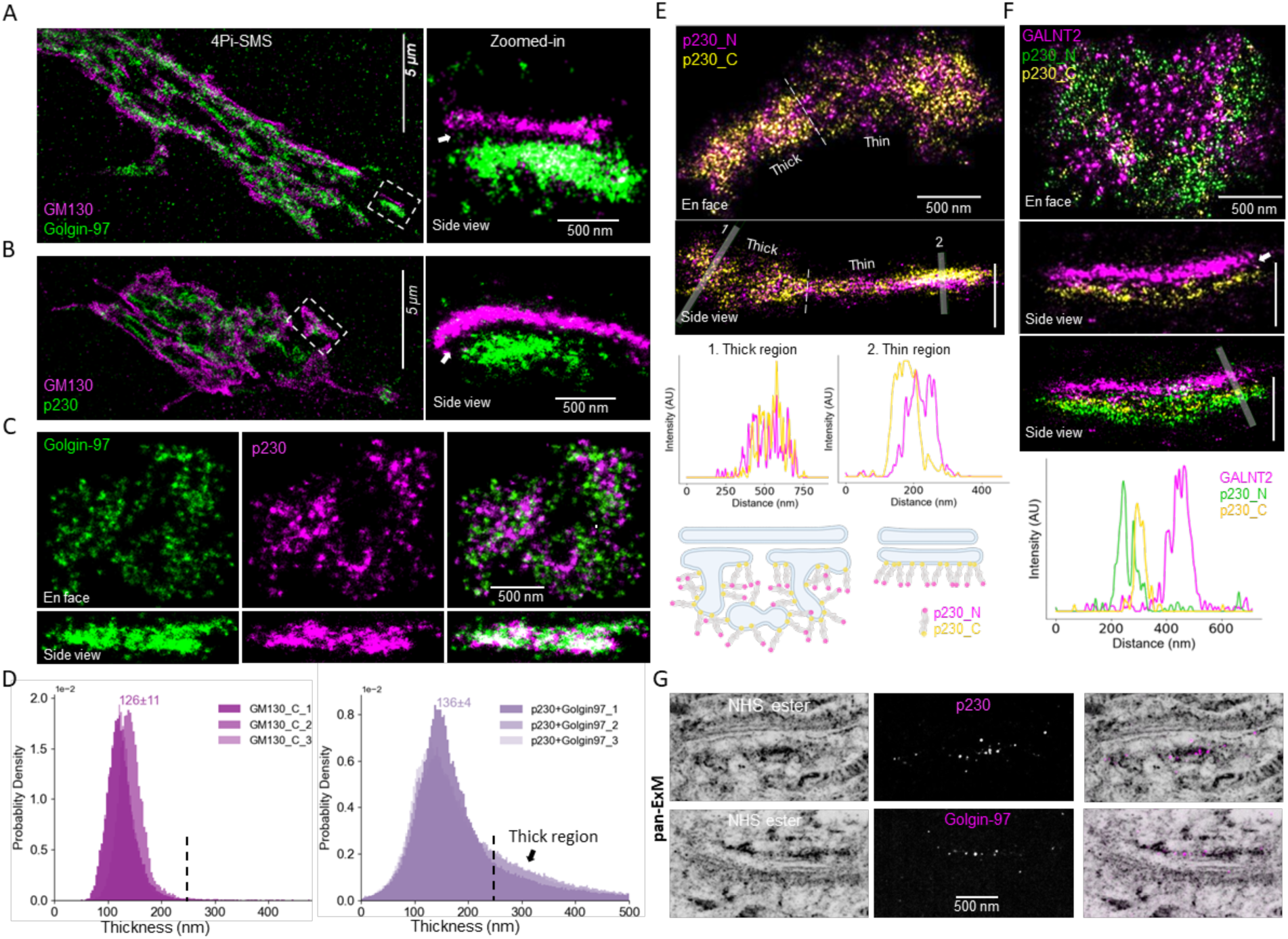
*Trans* cisternae have thin and thick regions. A. Two-color images of Golgin-97 with GM130. Left: whole cell. Right: Side view of cropped region (white box in Left). Arrow denotes the gap in-between. B. Two-color images of p230 with GM130. Left: whole cell. Right: Side view of cropped region (white box in Left). Arrow denotes the gap in-between. C. Zoomed-in images of p230 with Golgin-97 in the same cell. D. Isosurface thickness histograms of GM130 (Left) and p230 + Golgin-97 (Right). E. Two-color images of antibodies targeting p230_N (1-150 AA, rabbit) and p230_C (2063-2179 AA, mouse) (Top). Dotted line shows the separation between thick and thin regions. Transparent white bands show the lines used for the line-scan plots (Middle). Bottom cartoons show the two scenarios of p230 organization on the *trans* membrane. F. Three-color images of GALNT2 (sheep), p230_N, p230_C. Transparent white band show the line used for line-scan plot. Arrow denotes the gap in-between. G. Pan-ExM images of p230 and Golgin-97 in the context of Golgi stacks.

To examine whether the thin regions truly consisted of one cisterna only, we simultaneously labeled both N and C termini of p230 with antibodies in the same cell. Human p230 is a golgin of 2230 AA length and is anchored to Golgi membranes by GRIP domains at its C-terminus. We identified both thick and thin regions of p230 even in the same cisterna (Fig. 4E, movie S5). Surprisingly, seen in a side view, the N and C ends of p230 were well-separated in the same thin region as shown by line profiles (Fig. 4E top and middle panels). This is consistent with the idea that the thin region consists of sheet-like domains within the *trans*-most cisterna, as illustrated (Fig. 4E bottom). If true, we would expect that p230_C (containing GRIP) is closer to the Golgi membrane surface than p230_N. Using 3-color 4Pi-SMS, we indeed found that GALNT2 was closer to P230_C than to p230-N (Fig. 4F). There was even a small gap between p230-C and GALNT2 (Fig. 4F middle, white arrow), indicating an unlabeled 4^th^ cisterna, consistent with our assignments of cisternae to Golgin Layers.

Within the thick (about 250-500 nm) region of the TGN, however, the N and C ends of p230 are not resolved (Fig. 4E). The thick region is not likely to consist simply of two flattened and typical Golgi cisternae because (i) it is much thicker than the other two-cisternae golgins like GCCs (Fig. 2J) and (ii) we do not see a second peak in the SDF histogram (Fig. 4D). We suggest instead that the thick region corresponds to the topologically complex mixture of membranes including vesicles, tubules and tubular networks which has been well-described by EM of the TGN (De Matteis & Luini, 2008; Engel et al., 2015; Rambourg & Clermont, 1997). Notably, the *trans* sheet golgins are absent from the local rims (Fig. S7C), similar to the separation between the *cis* sheet golgin GM130 on the extended *cis* face being well-separated from Giantin, which is found only at the rim (Fig. 1B). In a side view, p230 extends beyond Layer 4 (marked by rim golgin GCC185) (Fig. S7C, D) as expected for the known tubule-vesicular regions of the extended TGN. Finally, we used pan-ExM to visualize p230 and Golgin-97 in the context of all the Golgi cisternae. As expected, both were located only at the *trans*-most region of Golgi stack (Fig. 4G).

### Sheet golgins form exclusively parallel dimers that polymerize into irregular meshes

Based on their localization within the Golgi, the known golgins fall into two clear categories, those located in four distinct layers at the curved outer rims (GMAP-210, Giantin, CASP, TMF, Golgin-84, GCC88 and GCC185) and those found on the enclosed membranes in sheet-like domains at the extreme *cis* (GM130) and *trans* (Golgin-97 and p230) faces of the stack. In light of our overall hypothesis that golgin self-assembly contributes to the organization of the Golgi (Rothman, 2019), it can be expected that the intrinsic physical chemistry of golgin proteins could reflect their apparently distinct preferences for the rim vs. sheet surfaces.

To investigate this, we comprehensively expressed and purified 7 full-length rim and sheet golgins and characterized their biophysical properties (Fig. 5AB). Only the terminal regions of GMAP-210 have previously been studied, and these were found to tether to flat or curved membranes, respectively (Drin et al., 2008). Full length GCC185 (Cheung et al., 2015) was previously purified and characterized and found to assemble into dimers with extensive rod-like regions separated by linkers whose flexibility is important for their function as tethers (Cheung et al., 2015).

**Figure 5.**
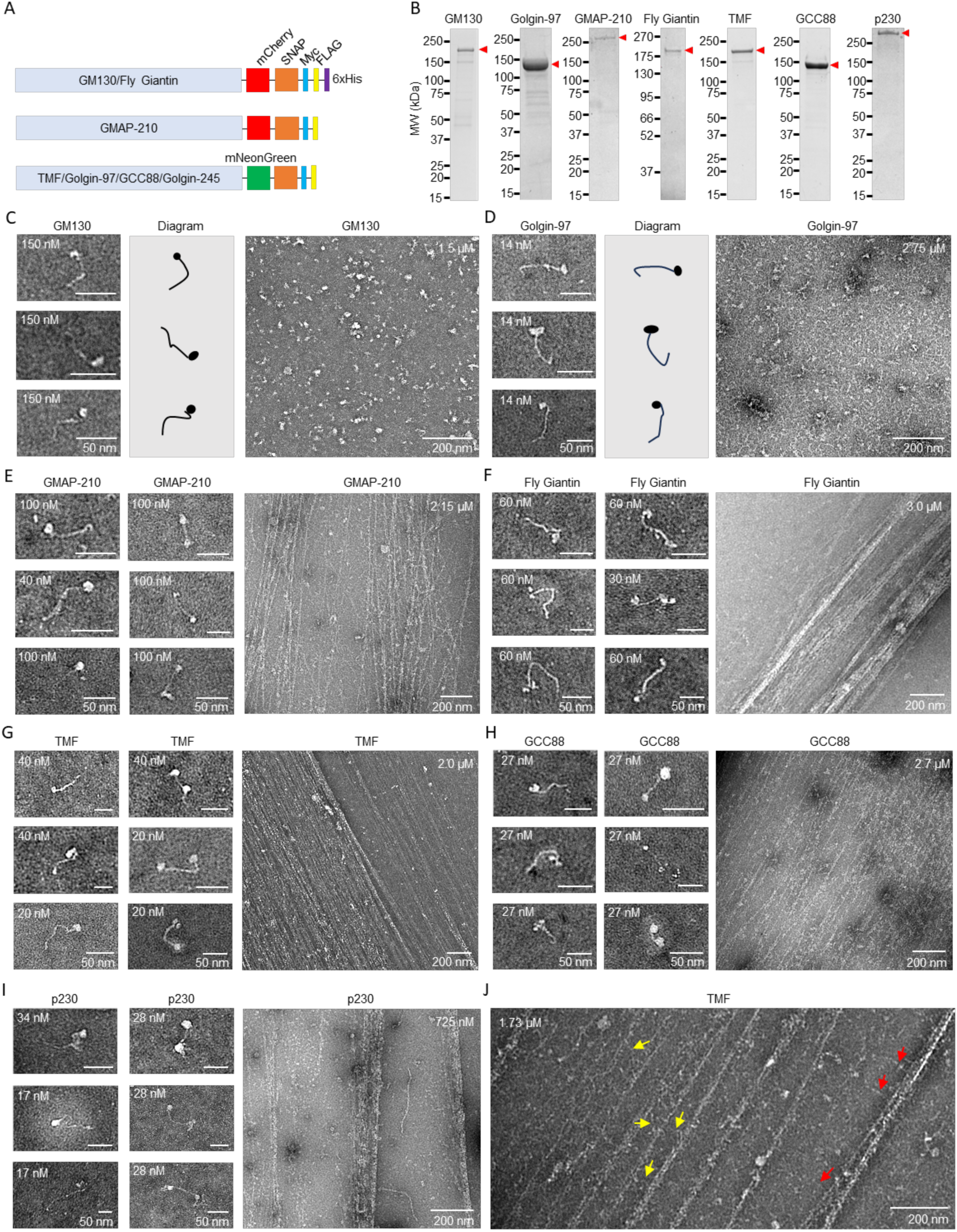
Structural analysis of purified golgins using negative stain electron microscopy revealed distinct organizational behavior for sheet and rim golgins. A. Domain diagrams detailing the position of C-terminal bulky tags on the constructs used for the expression and purification of selected golgins. B. SDS-PAGE of the elution fractions of the golgins purified by affinity chromatography using FLAG tag. Red arrowheads indicate the correct molecular weight for each protein. C-D. Negative stain EM images of purified GM130 and Golgin-97 at low (Left panel) and high concentrations (Right panel). Cartoon represents the parallel oligomers (Middle panel). E-I. Negative stain EM images of purified GMAP-210, melon fly Giantin, TMF, GCC88 and p230 respectively at nanomolar concentration range show the presence of parallel (Left panel) and anti-parallel oligomers (Middle panel). At micromolar concentration range, formation of bands of long filaments were observed (Right panel). J. Magnified negative stain EM image of TMF filaments. Red arrows indicate the rod-like arms that extend radially and irregularly from filaments. Yellow arrows indicate some of these rod-like arms that link filaments to each other.

The purified sheet golgins GM130 and Golgin-97 of *cis* and *trans* Golgi cisternae, respectively, both consist of rod-shaped molecules with a bulky spherical head of 5-7 nm diameter at one end when examined at 10-150 nM concentrations (Fig. 5C, D, Left, and S9). The length of the rod portion ranged from 50-60 nm for GM130 and 50-80 nm for Golgin-97 respectively. The spherical head results from mCherry/mNeonGreen and SNAP tags that were added at the C-terminus of these proteins in order to identify their orientation in EM analysis, as the C-terminus represents the primary membrane-attachment site of each golgin (Witkos & Lowe, 2015). Mass photometry measurements of GM130 and Golgin-97 revealed that dimers were the highest oligomeric species at concentrations between 10-100 nM, without clear peaks corresponding to higher oligomers. (Fig. S8). The two subunits in GM130 and Golgin-97 dimers have a parallel orientation, as the globular heads are never seen at both ends of a rod. We note that GM130 was previously reported to be a parallel tetramer (Ishida et al., 2015). This may be a higher association state present at a higher concentration. Consistent with this, although there was no clear tetramer peak, GM130 showed in consistently more frequent detection events in the tetramer range than other golgins (Fig. S8). Notably, both sheet golgins had a strong propensity to self-associate at concentrations > 1 μM (Fig. 5CD, Right, and S9) as irregular mesh-like aggregates, consistent with condensate formation noted in earlier work (Rebane et al., 2020). In contrast to rim golgins (next section) regular filaments were never observed.

### Rim golgins form mixtures of parallel and anti-parallel dimers that polymerize into cross-linked bands of multi-micron long filaments

GMAP-210 (Layer 1), Giantin (Layers 2+3), TMF (Layer 3), and GCC88 (Layer 4) were selected as representative rim Golgins to investigate. We expressed and purified human clones for these selected golgins except Giantin for which we used the more tractable melon fly orthologue because of its smaller molecular weight (120 kDa vs 376 kDa of human Giantin) and the absence of a trans-membrane domain (Fig. 5AB).

Mass photometry measurements of rim golgins (10-100 nM) revealed dimers as the highest oligomeric species without any peaks corresponding to monomers or higher order oligomers (Fig. S8). Negative stain EM images of the various rim golgins (10-100 nM) revealed two distinct populations of rod-shaped molecules with globular heads. In one case, the head was present only at one end of the rod (resembling the parallel dimers formed by GM130 and Golgin-97) (Fig. 5E-I, Left). In the second case, a head was present at both ends of the rod portion (Fig. 5E-I, Middle). Because only dimers are present, it is likely that the first population consists of parallel dimers (each head containing 2 unresolved mCherry/mNeonGreen and SNAP domains) and that the second population consists of anti-parallel dimers.

When examined at 10-100X higher concentrations (0.5-3 μM) all of the rim golgins now further assembled into strikingly long, thin filaments (Fig. 5E-I, Right and S10). The diameters of the individual filaments were quite uniform, ranging between 3-5 nm. However, the length of the filaments was highly variable, ranging from 1 to frequently >20 µm (often apparently limited by the edges of the EM grid). Although such elongated filaments were occasionally single, most filaments were further organized into ribbon-like “bands” which appeared to result from length-wise association of parallel filaments. Numerous linear rod-like arms (red arrows in Fig. 5J) extended radially and irregularly from the filaments and many but not all (yellow arrows in Fig. 5J) appeared to link the filaments to each other to assemble the bands. These arms are approximately the length of the individual rod-shaped dimers that predominate at 10-100 nM concentration. The width of the multi-micrometer long bands was variable, ranging up to ∼250 nm; but they always appeared to be very thin, likely comprised of a monolayer of laterally associated ∼3 nm single filaments. The bands as a whole tended to have distinct convex and concave sides, typically but not always being curved perpendicular to their long axis (Fig. S10, white arrows).

To test the hypothesis that the filaments and bands represent successive states of equilibrium association at high (μM) concentrations of the dimers observed at low (nM) concentrations, we chose TMF for a more detailed investigation. When first purified, TMF was present at 1 μM and had the form of bands. Dilution to 20 nM then resulted in only individual rods, previously shown to consist of a mixture of parallel and anti-parallel dimers. Then, to enable re-assembly, we concentrated the same sample back to 1.7 μM. As anticipated, the filaments and bands reappeared, confirming that they are assembled in a reversible process (Fig. 6A). We note that the bands that self-assemble from purified rim golgins can be sufficiently long (>20 µm) to form the rims that enclose much or even all Golgi apparatus. We also note that the width of these bands is compatible with the thickness of a Golgin Layer.

**Figure 6.**
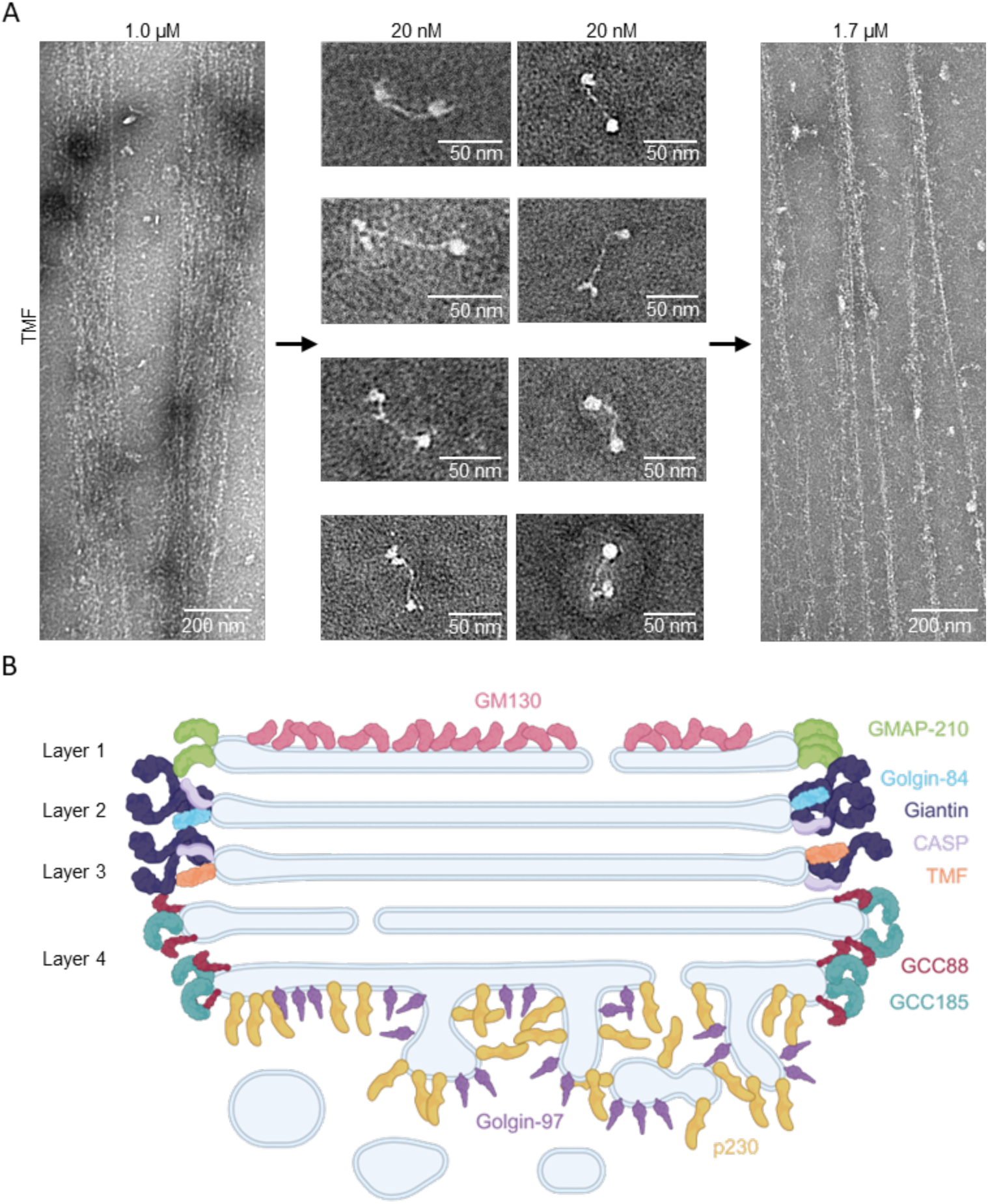
Filament formation by Rim golgins is concentration dependent and reversible. A. Negative stain EM images of TMF show that the filaments (Left panel) formed at micromolar concentrations (1.0 µM) fall apart to form anti-parallel rod-shaped molecules at 20 nM (Middle panels). When the diluted TMF was concentrated back to micromolar concentrations (1.7 µM), the long filaments reappeared (Right panel) indicating that filament formation is reversible. B. Schematic diagram of all the sheet and rim golgins imaged in this work. They all localize in the outer surface of Golgi apparatus.

Strikingly, purified p230, despite its localization to sheet-like domains at the *trans* face of the stack, nonetheless self-assembled into filaments and bands at low μM concentrations (Fig. 5I). This suggests that p230 has the potential to be a rim golgin in addition to its typical form *in situ* as a sheet golgin.

## Discussion

### Four compositionally distinct layers of Golgins

In this study we have comprehensively visualized the 3D organization of golgin proteins in the whole Golgi apparatus with unprecedented resolution at the 10-20 nm level, i.e. on the scale of the golgin molecules themselves. Remarkably, the outer perimeter of the Golgi ribbon was found to contain 4 distinct layers of golgins, each with a unique molecular composition. It is notable that the distribution of rim golgins amongst the 4 layers is precise at the single-cisterna level. That is, each of the first 3 membrane cisternae contains a unique combination of golgins at its rim, as do later cisternae as a group. The two glycol-processing enzymes tested in this paper also follow a precise discrete distribution. By contrast, previous reports described golgins, processing enzymes, Rabs, and SNAREs residing in the cisternae typically distributed in overlapping gradients spreading across multiple cisternae (Cosson et al., 2002; Gilchrist et al., 2006; Orci et al., 2000; Rabouille et al., 1995), more akin to peaks in a chromatography (or distillation) process (Glick et al., 1997; Rothman, 1981). A detailed comparison of our results with previous immuno-EM and Airyscan microscopy is provided in Table S4.

Utilizing proteomics data available for HeLa cells (Itzhak et al., 2016) we can estimate both the local concentration of individual golgins within a Golgin Layer and the approximate protein chemical composition of each Layer as follows: Assuming the vast majority of the rim golgins are Golgi-localized, Layer 1 appears to consist of GMAP-210 as the sole rim golgin (about 79,000 copies/cell). Layers 2 and 3 each contain Giantin (about 208,000 copies/cell) and CASP (126,000 copies) in approximately equal amounts based on relative staining (see Fig. 2A-D and S6A for examples), suggesting that each Layer contains about half (104,000 copies) of Giantin and about half (63,000 copies) of CASP. However, Layer 2 also uniquely contains Golgin-84 (about 48,000 copies) and Layer 3 uniquely contains TMF (about 43,000 copies). This suggests that Layer 2 consists of Giantin, CASP, and Golgin-84 in the rough proportions of 2:1:1; and that Layer 3 consists of Giantin:CASP:TMF in similar 2:1:1 molar proportion. Layer 4 contains only GCC88 (about 69,000 copies) and GCC185 (about 99,000 copies), roughly in a 1:1 ratio. Because we have localized all of the canonical rim golgins, we can reasonably assert that there are no other golgins within these four rim Layers.

### How do filaments of rim golgin assemble into bands?

From our biochemical experiments, we noted that all rim golgins assemble into rod-like dimers. Because of their characteristic abundance of sequences having a high propensity to form rod-like α-helix-based coiled-coils, it is natural to assume that the rods are composed of two-stranded coiled-coils, whose orientation can in principle be either parallel or anti-parallel. Based on the localization(s) of protein tag(s) within the dimers, we concluded that both isomers are present as a mixture (Fig. 5). This suggests simple models for how filaments could self-assemble from dimers and be flexibly (and potentially fluidly) cross-linked into bands of filaments. For example, the anti-parallel dimers could assemble into the extensive linear filaments, while parallel dimers could cross-link the filaments. This would explain why rim golgins can form cross-linked filaments, but sheet golgins can only form cross-links (i.e., meshes). In one way or another, cross-linking implies that the coiled-coil regions within the golgins are unstable and capable of alternative intra- and inter-molecular arrangements. Numerous examples of instability and alternative conformations have been highlighted in earlier studies of single golgin molecules (Cheung et al., 2015; Drin et al., 2008; Ishida et al., 2015).

Many arrangements are possible of course and more detailed structural and biochemical studies will be needed to differentiate among models. Because characteristic mixtures of rim golgins compose each Layer, it will be interesting in future research to see if corresponding mixtures of purified golgins form hybrid bands and if so how.

### Is the golgin tetraplex observed in cells composed of aligned bands of filaments?

Is it likely that the golgin proteins are assembled into the filaments and bands within the Golgi apparatus? We noted that filaments (laterally associated into bands) prevail *in vitro* at concentrations typically above 1 μM. We can estimate the local concentration of golgins at the surface of the Golgi apparatus using the same approach as in previous work (Rebane et al., 2020). The results vary from ∼15 μM (TMF and Golgin-84) to ∼60 μM (Giantin) easily in the range at which filaments and bands predominate *in vitro*. Given the extreme lengths of the filaments, which can even exceed the entire Golgi contour in length, it is expected that the filaments are circumferentially oriented such that the bands they form will enclose and possibly shape the rims of stacked membrane cisternae they surround. Future studies using cryo-electron tomography *in situ* will be needed to test these ideas. It is worth noting that some intracisternal filament bundles can be observed within trans-Golgi buds (Engel et al., 2015).

While here we are emphasizing novel physical roles of collective assemblies of golgin filaments and bands, it is important to recall the well-established biochemical roles of these same proteins (functioning individually) in capturing transport vesicles and bringing them to the surface of the enclosed Golgi membranes for fusion (Lowe, 2019). How can the same protein do both? The obvious possibility is that, while many copies of the rim golgin form a stable casement stabilizing the Golgi ribbon, other copies are simultaneously available as tethers reaching out from the casement. It is easy to imagine a dynamic equilibrium in which anti-parallel dimers alternate between two states: cross-linking two filaments in the casement in one state; and with one end free to capture a vesicle in the other state. This dynamic equilibrium would enable specific classes of transport vesicles (bearing unique combinations of Rab[GTP] proteins) to be captured at the outer surface of each rim layer by the unique combination of golgins comprising the casement in that layer. The vesicles would then dissolve the casement locally through further multivalent Rab interactions, essentially diffusing across in a facilitated manner not unlike the manner in which selected cargo traverses nuclear pores (Rothman, 2019).

Altogether our data suggest a simple and novel “outside-in” view of the architecture of the Golgi, in which four distinct bands of golgins stack on each other to define the shape and potentially the content of membranes within the organelle (Fig. 6B). Given their dual role as specific vesicle tethers, each Golgin Layer could import cognate membrane content allowing selective access for fusion within (Rothman, 2019). In the next section we will suggest how the golgin tetraplex could organize the cisternal compartments from the outside-in, be it in interphase or in Golgi re-assembly following cell division.

### The Golgin Organizer Hypothesis, a physical model for how the tetraplex Golgi rim could self-organize and result in a stack of enclosed membrane cisternae

Basic physical considerations suggest how rigid protein filaments could stretch and flatten attached condensates to yield a disk (Graham et al., 2023). Similar mechanisms may exist for rim golgin filamentous bands that attach to Golgi membranes. We postulate that the four golgin bands adhere to each other in the order in which they are observed *in situ*. We noted that the bands forming *in vitro* typically have a concave and a convex surface. We suggest that it is the concave surface that binds the Golgi rim. Then, in order for this casement to generate a stack of flat cisternae within, there must be a strong preference for binding highly curved bilayers at the inner (concave) surface of each band. This curvature binding preference is already well-known for GMAP-210, whose dimeric ALPS domains bind preferentially to highly curved bilayers (Drin et al., 2008). Other rim golgins have been noted to have ALPS-like domains (Wong et al., 2017) or are attached by TMDs or combinations of bilayer-inserted prenyl and fatty acyl chains linked to reversibly associated GTPases, which may ultimately prove to have a curvature preference.

Given the above, so long as the total favorable energy gained from golgin curvature-binding domains exceeds the unfavorable energy required to bend and flatten an enclosed vesicle, each Golgin Layer will deform such a vesicle into an enclosed flat cisterna such as are observed in the Golgi stack. Whatever the details, the overall energy will also be minimized by reducing the overall curvature of the rim in the plane of the Golgi ribbon. This favors coalescence of ministacks into the ribbon and will ultimately result in a single ribbon in every cell. To the extent that the sheet golgins at the opposite *cis* and *trans* faces of the stack prefer flat over curved membranes, their membrane adhesion will synergize with the curvature-inducing rim golgins to stabilize a stacked disc morphology of membranes within the Golgin-encased ribbon.

In our speculative physical hypothesis, Golgi membrane morphology is generated solely from the “outside-in” by adhesion of the outer surface of the Golgi membrane system to a casement of self-organizing golgin proteins. This differs radically from traditional “inside-out” models in which the cisternal membranes are glued together internally by hypothetical “stacking proteins”. Despite decades of research no such proteins have yet been found. Various candidates have been proposed, notably GRASP proteins, based on compelling *in vitro* reconstitution experiments (Barr et al., 1997; Shorter et al., 1999; Wang et al., 2003) and notably also Golgin-45 based on Golgi morphological changes in knockdown cells (Lee et al., 2014; Short et al., 2001), but their effects can now be alternatively explained physically as “outside-in” by stabilizing the adhesion of GM130 sheets to the *cis* face. Indeed, deletion of both GRASP genes had no effect on the stacking *in vivo* (Grond et al., 2020; X. Zhang & Wang, 2020). It is also important to point out that in this comprehensive study we report that there are no golgins present anatomically in the inter-cisternal spaces where “stacking proteins” would need to exist. Indeed, the key feature of our hypothesis is that no “stacking proteins” are needed. The rim golgins would function as the proposed curvature stabilizers as required for *in silico* generation of Golgi-like morphologies (Tachikawa & Mochizuki, 2017).

Our physical model presumes that the Golgi membrane vesicles have ion permeabilities and/or pumps that enable or favor deflation, as indicated by classic experiments in which various ionophores cause the cisternae to swell (A. Tartakoff et al., 1978; A. M. Tartakoff, 1983). Indeed, these long-puzzling observations are now readily explained.

Genetic studies are also broadly consistent with the Golgin Organizer hypothesis. The otherwise puzzling and mild effects of golgin gene knockdowns and knockouts (Witkos & Lowe, 2015) can now easily be rationalized. Typically, these are not lethal, and secretion through the Golgi continues, but the morphological effects are complex and seem hard to explain. Accompanying this, the single Golgi ribbon (“Golgi area”) typically fragments into smaller ribbons or aggregates of ministacks, or even completely into ministacks, especially when more than one golgin is removed at a time. Our model predicts this continuum of behaviors, because it is the *total* energy that determines morphology so that no *single* golgin would be critical.

### Limitations of this study

There are several limitations of the current work. Because we focused on endogenous levels, our results were restricted by the availability of specific and efficient primary antibodies from limited species, which may not exist for all the Golgi-resident proteins and their specific regions. All used antibodies in this work are listed in Table S1. Golgin-160 was not included in this study but is the subject of a different project. We did not systematically address cell-to-cell variation in the numbers of cisternae and how it affected golgin localization due to relatively low throughput of super-resolution and EM imaging. Finally, this observational study does not rule out other possible mechanisms underlying the targeting of golgins to specific compartments, and parallel mechanisms of GA organization by other Golgi-resident proteins. Future detailed genetic and biochemical studies are necessary to establish causal relationships between golgin localization and Golgi architecture.

## Acknowledgements

We thank Alexandra Louka for cloning and purifying melon fly Giantin, Zach Marin and Lukas Fuentes for help with PYMEVisualize, Yale School of Medicine CCMI for confocal and electron microscopy, Frederic Pincet, Vivek Malhotra, Ishier Raote, Iván López Montero, Min Wu, Anup Parchure for discussions. J.B. acknowledges support by the Wellcome Leap Foundation and the National Institute of General Medical Sciences (R01GM151829).

## Author contributions

M.S. and J.E.R. conceived this project. M.S., A.R. and. J.E.R. designed the research. M.S. performed 4Pi-SMS experiments. A.R. performed negative stain EM experiments. Y.Y. designed and performed isosurface-based thickness analysis. Y.T. and O.M. performed pan-ExM experiments. M.G. and X.L. performed immuno-EM experiments. J.N.D.G. and A.R. performed the mass photometry experiments. J.C. made plasmids and H.Z. purified all the proteins. Y.Z. contributed to 4Pi-SMS methodology. Y.Y. and M.S. drew illustrations. M.S., A.R., J.B. and. J.E.R. wrote the manuscript with input from all authors. J.E.R. and J.B. supervised the study.

