## Supplementary material for "The Golgi Rim is a Precise Tetraplex of Golgin Proteins that Can Self-Assemble into Filamentous Bands": Methods and Supp Figures and Tables

#### Cell culture

HeLa CCL2 (ATCC) were cultured in DMEM (Gibco) with 10% FBS (Gibco) at 37°C with 5% $CO_2$. Pooled HUVEC (Lonza) were cultured in in EGM-2 Endothelial Cell Growth Medium (Lonza). SAOS-2 HTB-85 (ATCC) were cultured in McCoy's 5A medium (Cytiva) with 15% FBS (Gibco). Transfection of ManII-GFP was performed by electroporation using a Neon™ Transfection System according to manufacturer's protocol (Invitrogen).

#### Plasmids

ManII-GFP (amino acid 1-88 of rat alpha-mannosidase 2) was cloned into pEGFP-N1 to have C-terminal eGFP in the Golgi lumen. Human golgin genes with C-terminal Myc-FLAG tags were purchased from Origene: GM130 (RC209641), GMAP-210 (RC220674), TMF (RC220590), GCC88 (RC204339), Golgin-245/p230 (RC230747). The Golgin-97 gene (RG206064) was purchased with TurboGFP. Melon fly giantin (Uniprot ID: A0A0A1XP57) was synthesized (GeneScript) with mCherry-SNAP-Myc-FLAG-6xHis C-terminal tags. GMAP-210 (G1827S variant), TMF, GCC88 and Golgin-245/p230 were altered by inserting a PCR fragment containing either mCherry or mNeonGreen linked to a SNAP tag at the sole NotI restriction site located before the myc tag using In-Fusion Cloning (Takara Bio). For GM130 and Golgin-97, the genes were amplified by PCR and cloned into a pClneo expression vector (Promega) that contains either mCherry-SNAP-Myc-FLAG-6xHis or mNeonGreen-SNAP-Myc-FLAG C-terminal tags, respectively.

#### Protein purification

Golgin DNA constructs were transfected into Expi293 cells (ThermoFisher) according to the manufacturer's instructions and expressed for up to 48 hours. Proteins were purified using C-terminal FLAG tag as described in our previous publication(Rebane et al., 2020), with some minor modifications. To summarize, cell pellets from 50ml cultures were resuspended in 10ml of buffer [50 mM HEPES pH7.3 (Gibco), 175mM NaCl, 1mM TCEP, 10ug/ml DNaseI, 1mM PMSF, and proteinase inhibitors (Roche)]. Cells were dounced, the volume was increased to 20ml and then rotated at 4C for 20 min. The lysate was centrifuged at 17500xg for 30 min at 12C in a Beckman JA-20 rotor. The supernatant was removed and 0.6ml of prewashed FLAG resin slurry (Sigma) was added and allowed to bind for 2 hours at 4C. Afterwards, the resin was moved to a column (BioRad) and washed with 20ml of buffer (omitting the DNase and PMSF) followed by an ATP treatment of buffer supplemented with 1mM ATP and 1mM $MgCl_2$ for 30 min at room temperature. The resin was then washed with 20ml of buffer, 10ml of buffer plus 5mM EDTA, and finally with the buffer supplemented with 1mM EDTA, 5% glycerol, and 125ng/ml 3X FLAG peptide (APExBIO) to elute the protein. Elutions were done in succession of 0.55ml, 0.45ml, and 0.35ml for 1 hour, 45min and overnight, respectively.

#### Primary and secondary antibodies

See Table S1 and S2 for the antibodies used in this work.

### Sample labeling for 4Pi-SMS microscopy

HeLa, SAOS2 or HUVEC cells were seeded on 25 mm or 30 mm diameter No. 1.5H round coverslips (Thorlabs, NJ) housed in a 6-well plate. They grew for 1-3 days before fixation by 4% paraformaldehyde (EMS, PA) in 1.5X PHEM buffer (EMS, PA) for 15 mins. Then samples were permeabilized with 0.3% NP40 + 0.05% TX-100 and blocked with normal serum (Jackson ImmunoResearch, PA) from the species of the secondary antibodies. Primary antibodies at 1:200-1000 were incubated overnight at 4 °C. Secondary antibodies were used at 1:500 dilution for 2 hours at room temperature. Samples were post-fixed in 3% PFA + 0.1% GA for 10 min, added 0.1 µm crimson beads (Invitrogen), and stored in PBS at 4 °C.

### 4Pi-SMS microscopy imaging and data processing

A custom-built 4Pi-SMS microscope was used for this study (Huang et al., 2016; Zhang et al., 2020). Sample mounting, image acquisition, and data processing were done similarly to those previously described, except for the imaging speed set at 200 Hz and a 642 nm laser intensity of ~12.5 kW/cm$^2$. Typically, one acquisition involved recording 100-200 imaging stacks, each comprising 3000 frames per stack, with 1-6 Z-steps of 500 nm step size. Raw images were processed using custom MATLAB code to generate localization data. Drift correction was performed using a minimum entropy method (Cnossen et al., 2021). The localization data were then imported into Vutara SRX 7.0.06 software (Bruker, Germany). For visualization purposes, all 4Pi-SMS images and videos were rendered using the Point Splatting mode (10-15 nm particle size). Some data sets were denoised based on local density using build-in function in Vutara SRX (Fig. S3C). Line profiles were generated using a custom ImageJ Macro code and plotted with either MATLAB or Python and Matplotlib package. Curve fitting (Gaussian) was performed with sci-kit package. Illustrations were generated in Biorender.com.

### Isosurface generation and thickness analysis

**Isosurface Generation:** 3D isosurfaces of the Golgi were created in PYMEVisualize (v23.05.17) using octree (Marin et al., 2021) and shrinkwrapping algorithm (Marin et al., 2023) (Fig. S5A). The parameters of the isosurface creation and shrinkwrapping algorithms were adjusted such that the 3D isosurfaces encompass the imaged localizations closely. **Data Cleaning:** The 3D isosurface meshes were then processed using MeshLab (v2023.12) (Cignoni et al., 2008). Small, disconnected components were removed from the surface meshes. Faces in the meshes were inverted for later calculation. Duplicated vertices were removed. Absolute curvatures of the vertices were calculated and used to filter out vertices both at the outer rim of the cisternae and inner rim of fenestration (Fig. S5C). **Measurement:** To estimate the thickness of the membrane mesh, we use shape diameter function (Shapira et al., 2008), which calculates the weighted average of a cone of rays sent from each vertex to the opposite side of the mesh (Fig. S5B). We choose 25 degrees as the cone amplitude to ensure the SDF values are robust and approximate to the actual thickness of the isosurfaces. Therefore, it can be proved that the SDF values should be between 1 to $\frac{1}{cos12.5°} \approx 1.024$ times of the actual thickness of the 3D isosurface meshes. The SDF value histograms (Fig. S5D) were generated using Python 3.10 with NumPy (v1.23.4), SciPy (v1.9.3), plyfile (v0.7.4) and matplotlib (v3.6.2) packages.

### pan-Staining expansion microscopy (pan-ExM)

pan-ExM was performed as previously reported (M'Saad & Bewersdorf, 2020). Briefly, Hela cells were fixed with 4% formaldehyde (FA) (EMS, 15710) in 1× PBS for 1 h at RT and then rinsed three times with 1× PBS. Next, samples were post-fixed with 0.7% FA and 1% (w/v) acrylamide (AAm; Sigma-Aldrich, A4058) in 1× PBS for 6 h at 37 °C. After washing the samples with 1× PBS, samples were embedded in the first expansion gel solution composed of 19% (w/v) sodium acrylate (SA; Santa Cruz Biotechnologies, SC-236893C), 10% (w/v) AAm, 0.1% (w/v) N,N'-(1,2-dihydroxyethylene) bisacrylamide (DHEBA; Santa Cruz Biotechnologies, SC-215503), 0.25% N,N,N',N-tetramethylethylenediamine (TEMED; American Bio, AB02020) and 0.25% (w/v) ammonium persulfate (APS; Sigma-Aldrich, A3678) in 1× PBS. Gelation was performed for 1.5 h at 37°C in a humidified chamber. Then samples were transferred into the denaturation buffer containing 200 mM sodium dodecyl sulfate solution (SDS; Bio-rad, 1610418), 200 mM Sodium chloride (NaCl; J.T. Baker, 3624-01), 50 mM tris [hyroxymethyl] aminomethane (Tris; American Bio, AB02000) and incubated at 73 °C for 1 h. After expanding the samples in MilliQ water, gels were incubated in a neutral gel solution (10% (w/v) AAm + 0.05% (w/v) DHEBA + 0.05% (v/v) TEMED + 0.05% (w/v) APS in 1× PBS) twice for 20 min each at RT. Next, residual gel solution was removed with Kimwipes, then the gels were sandwiched between two pieces of coverslips and incubated at 37 °C for 1.5 h in a nitrogen-filled humidified chamber. Following rehydrating the gels with 1 × PBS, gels were incubated in a second gel solution composed of 19% (w/v) SA, 10% AAm (w/v), 0.1% (w/v) N,N′-methylenebis (acrylamide) (BIS; Sigma-Aldrich, M7279), 0.05% (v/v) TEMED and 0.05% (w/v) twice for 15 min each on ice. Next, residual gel solution was removed with Kimwipes, then the gels were sandwiched between two pieces of coverslips and incubated at 37 °C for 1.5 h in a nitrogen-filled humidified chamber. To dissolve DHEBA, gels were incubated in 200 mM Sodium hydroxide (NaOH; Brand-Nu Laboratories, PC5960S-1) for 1 h at RT and then washed with 1× PBS. Next, the gels were immunolabeled with primary antibodies (GM130, Proteintech, 11308-1-AP; Giantin, Sigma, HPA011555; GLANT2, Novus, AF7507-SP; GFP, Thermofisher, A11122) at 1:250 for 24 hours at 4°C, and secondary antibodies (goat anti-rabbit IgG Atto647N; Sigma-Aldrich, 40839; Donkey Anti-Sheep IgG CF633; Biotium, 20134-1mg) at 1:250 for 24 hours at 4°C, then pan-stained with NHS ester Atto532 (Sigma-Aldrich, 88793) or NHS ester CF568 (Biotium, 92131)dyes. Before imaging, gels were expanded in MilliQ water for fully expansion, then mounted on glass-bottom dishes (MatTek, P35G-1.5-20-C) and sealed with two-component silicone glue (Picodent Twinsil, Picodent, Wipperfürth, Germany). Samples were imaged on Leica SP8 STED 3X or Dragonfly High Speed Confocal Microscope Systems (Andor/Oxford instruments). Expansion factor was estimated to be ~15 (linear).

**Cryo immuno-electron microscopy**

Cell cultures were fixed in 4% paraformaldehyde + 0.2% glutaraldehyde in PBS for 30minutes followed by further fixation in 4% PFA for 1 hour. They were rinsed in PBS and re-suspended in 10% gelatin. The blocks were trimmed and placed in 2.3M sucrose overnight on a rotor at 4C and were transferred to aluminum pins and frozen rapidly in liquid nitrogen. The frozen blocks were cut on a Leica Cryo-EMUC6 UltraCut and 60 nm thick sections were collected using the Tokuyasu method (Tokuyasu, 1973) and placed on carbon/formvar coated grids and floated in a dish of PBS for immunolabeling.

Grids were placed section side down on drops of 0.1M ammonium chloride to quench untreated aldehyde groups, then blocked for nonspecific binding on 1% fish skin gelatin in PBS. The grids were incubated on a primary antibodies mouse anti-GM130 (BD) at 1:50. 10nm Protein A gold (Utrecht Medical Center) was used as a secondary antibody. All grids were rinsed in PBS, fixed using 1% glutaraldehyde for 5mins, rinsed, and transferred to a UA/methylcellulose drop before being collected and dried.

60nm sections on grids were viewed FEI Tecnai Biotwin TEM at 80 kV. Images were taken using AMT NanoSprint15 MK2 sCMOS camera.

**Negative stain electron microscopy**

The negative stain EM samples of golgin proteins were prepared directly from the eluted fractions at μM concentrations and from eluted fractions diluted to nanomolar concentrations using 25 mM HEPES 7.4, 140 mM KCl, 2 mM $MgAc_2$ and 1 mM TCEP. For preparing the negative stain EM grids, 5 μL of the protein samples were directly applied to a glow discharged Carbon Type-B, 400 mesh, Copper grid (Ted Pella). After 1-minute incubation on the grid, the samples were quickly washed with 5 μL of 2% (w/v) uranyl formate solution, followed by staining with another 5 μL of 2% uranyl formate for 1 minute. The grids were imaged using an 80 kV JEOL JEM-1400Plus microscope equipped with a bottom-mount 4k × 3k charge-coupled device camera (Advanced Microscopy Technologies). The images were processed and analyzed using the Gatan Microscopy Suite Software.

**Mass photometry**

For mass photometry measurements silicon gaskets attached to clean high-precision microscope cover glasses were mounted on the lenses of a mass photometer (Refeyn) covered with immersion oil. The holes in the gaskets were first filled with 10 μL of the sample buffer (25 mM HEPES 7.4, 140 mM KCl, 2 mM $MgAc_2$, 1 mM TCEP) for focusing the cover glasses. Once the focus is stabilized 5 μL of the golgin proteins were diluted from the stock using the sample buffer and added in appropriate amounts to get the desired final concentration in the 10-100 nM range. The data was acquired for 60 s using the AcquireMP software and analyzed using the DiscoverMP software. In all the cases there was a peak to the left of the dimer with an estimated molecule weight lower than the monomer, probably indicating impurities. We ignored it from our analysis. The contrast to mass calibration curve for the mass analysis was prepared each time using beta-amylase and thyroglobulin.

### Supplemental Figures

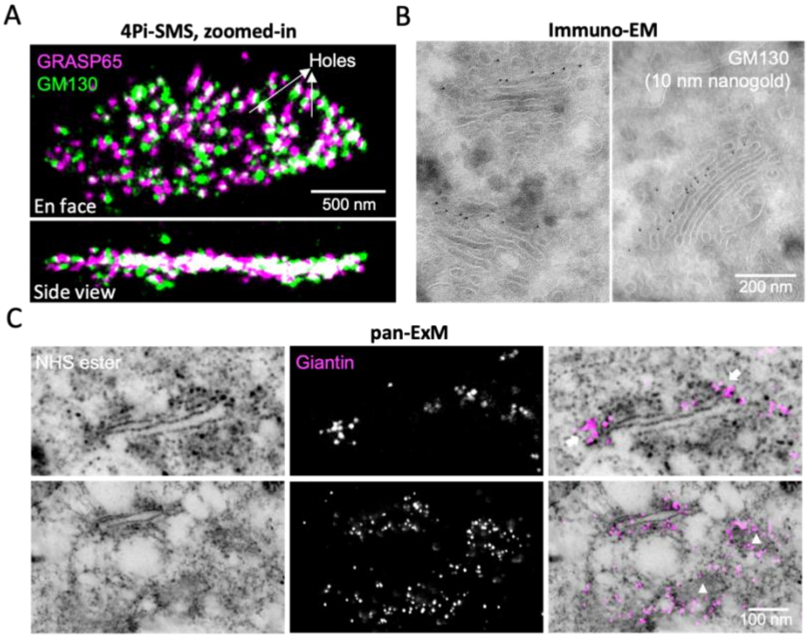


**Supplemental Figure 1. Sheet golgin GM130 and rim golgin Giantin.**

A. 4Pi-SMS image of GM130 and GRASP65 shows completely colocalization in both *en face* (Top) and side view (Bottom).

B. Additional EM images of immunogold labeled GM130.

C. Pan-ExM images of NHS ester (Left), Giantin (Middle) and merge (Right). White arrows show two sides of cisternae rims from cross-section of Golgi, and white triangles show en face of cisternae. Scale bars are corrected for the expansion factor.

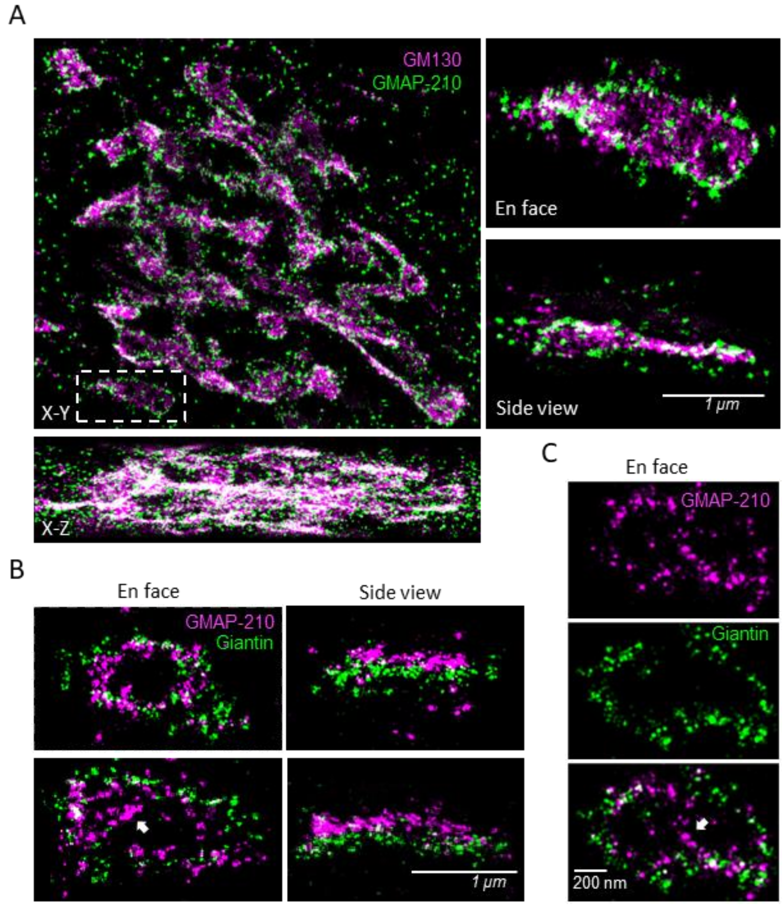


**Supplemental Figure 2. GMAP-210 locates at rim of *cis* cisterna.**

A. 4Pi-SMS images of GM130 and GMAP-210 (antibody against 1760-1855 aa). Right panels show a zoomed-in cisterna of white dashed box on the Left.

B. Additional 4Pi-SMS images of zoomed-in GMAP-210 and Giantin shown in both *en face* (Left) and side view (Right). White arrow shows the additional signal only in GMAP-210 channel.

C. Additional 4Pi-SMS image of zoomed-in GMAP-210 and Giantin shown in single color. White arrow shows the additional signal only in GMAP-210 channel.

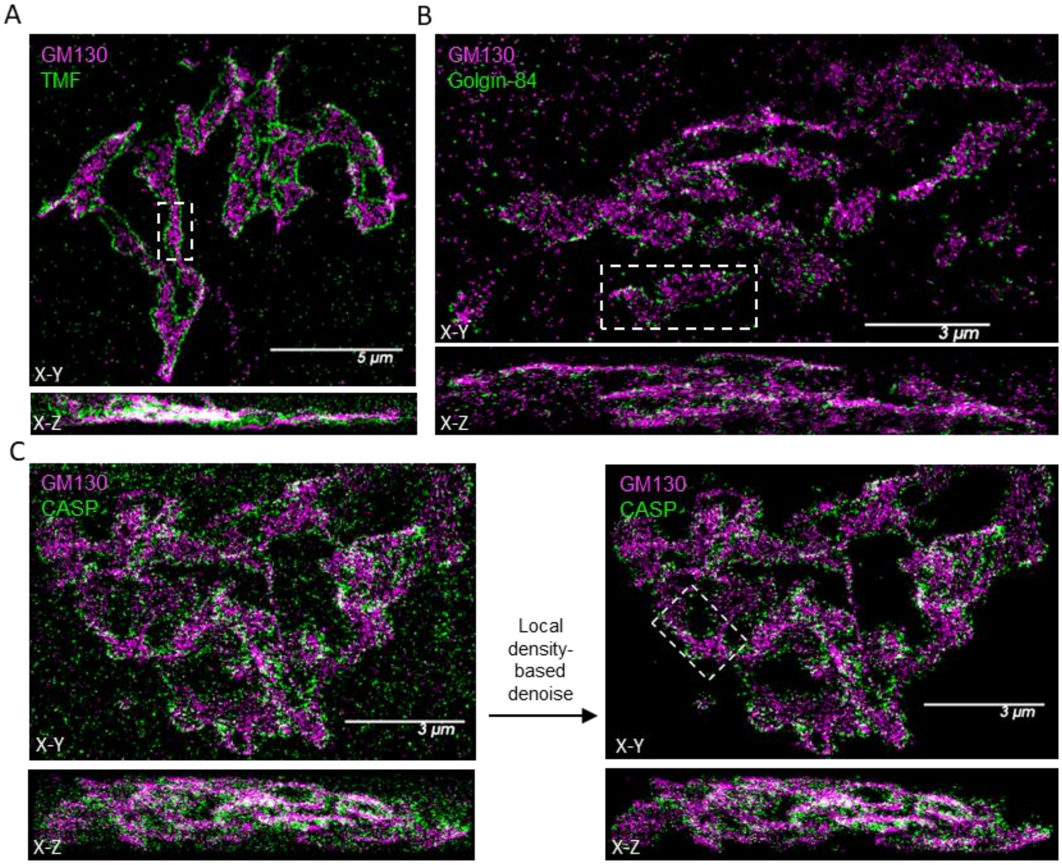


**Supplemental Figure 3. Overview of TMF, Golgin-84, CASP localizations at rims of medial cisternae.**

A-C. 4Pi-SMS overview images of the entire Golgi region in a cell labeled by GM130 and TMF (A), Golgin-84 (B) or CASP (C). The white dashed box shows the region used in Fig. 2A.

C. Local density-based denoised images before (Left) and after (Right).

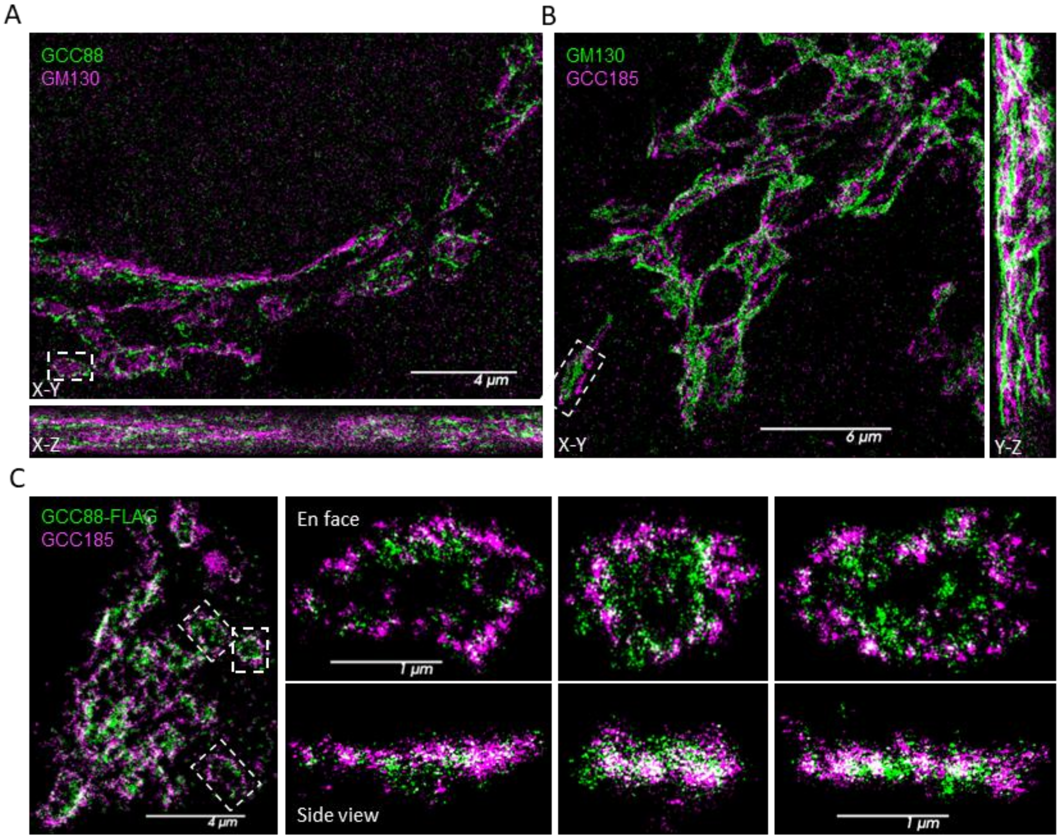


**Supplemental Figure 4. GCC88 and GCC185 locate at rim of *trans* cisternae.**

A-B. 4Pi-SMS overview images of the entire Golgi region in a cell labeled by GM130 and GCC88 (A) or GCC185 (B). The white dashed box shows the region used in Fig. 2A.

C. Overview images of overexpressed GCC-FLAG and endogenous GCC185 (Left) and three zoomed-in regions (Right) as outlined by white dashed boxes.

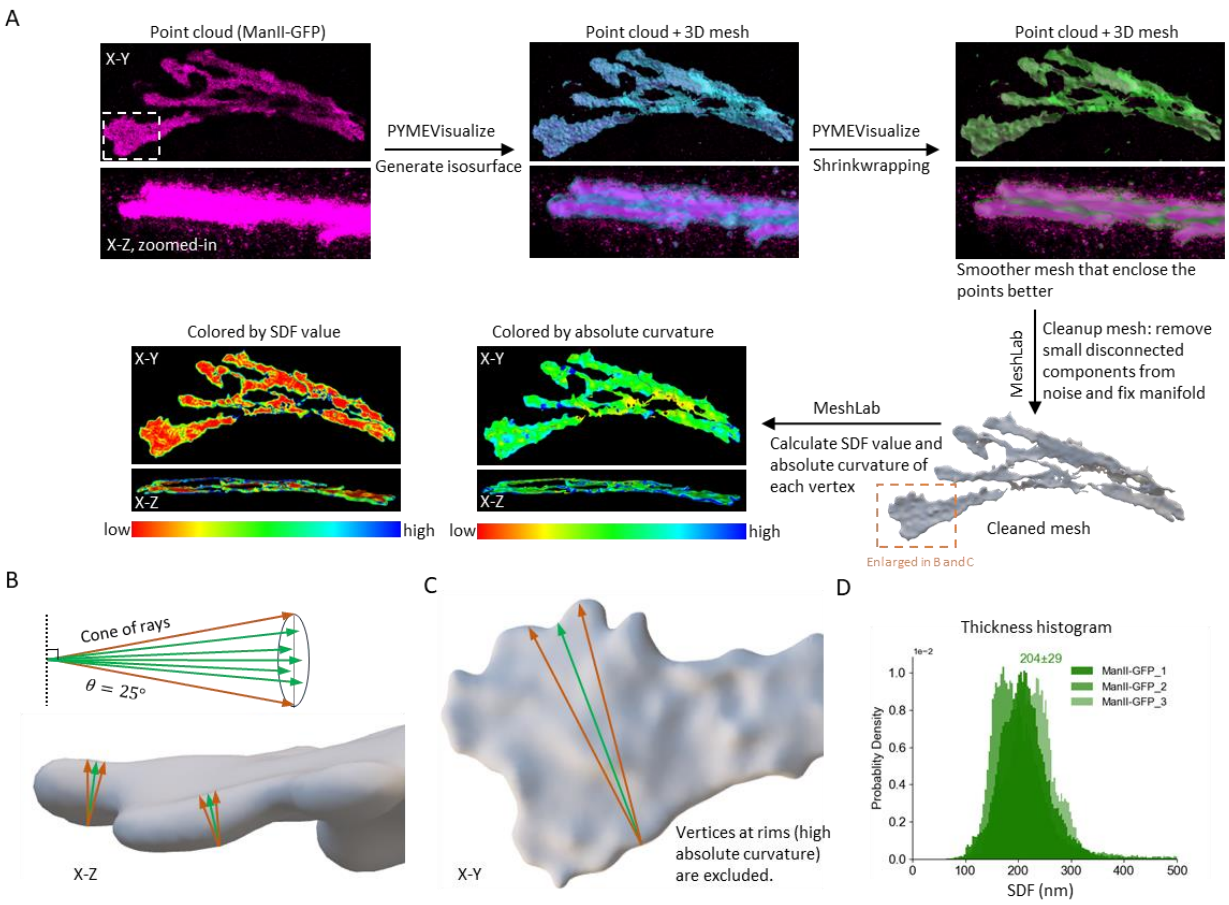


**Supplemental Figure 5. Procedure of isosurface generation and thickness analysis.**

A. Workflow of generating shrink-wrapped membrane isosurface meshes and computing SDF values. See also *Methods - Isosurface generation and thickness analysis*.

B. Illustration of the cones of rays sent from vertices on the sheet area (lower curvature part) of the isosurface mesh. These vertices are included for SDF computation.

C. Illustration of the cones of rays sent from vertices on the rim area (higher curvature part) of the isosurface mesh. These vertices are excluded in SDF computation.

D. A histogram plotted with the SDF values of all valid vertices of three ManII-GFP isosurface meshes. The SDF values at the peaks of the histogram represent the thickness of the isosurface meshes.

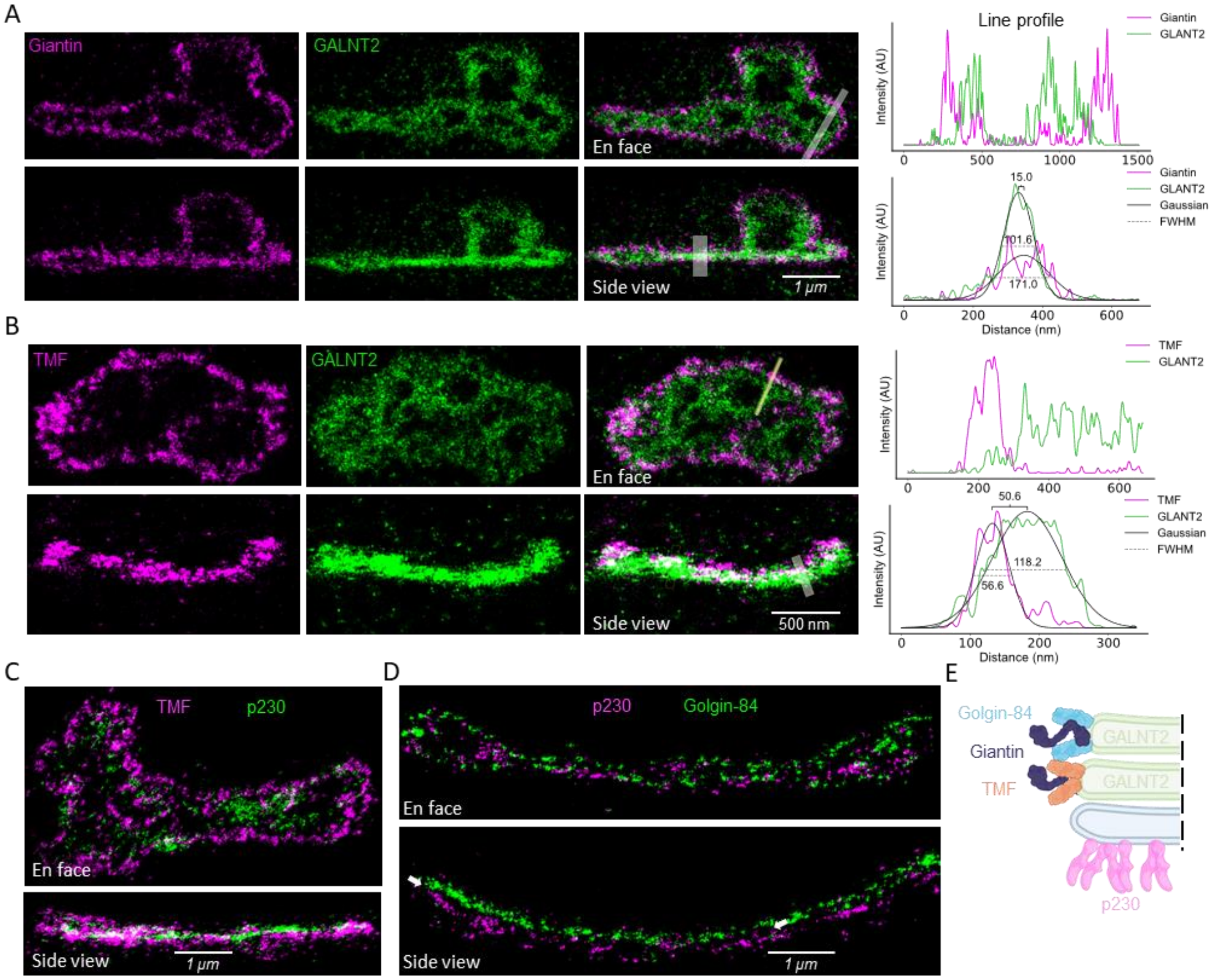


**Supplemental Figure 6. Relative positions between medial rim golgins and sheet GALNT2 / p230.**

A. 4Pi-SMS images of Giantin (magenta) and GALNT2 (green) with line-scan profiles (Right).

B. 4Pi-SMS images of TMF (magenta) and GALNT2 (green) with line-scan profiles (Right).

C. 4Pi-SMS images of TMF (magenta) and p230 (green).

D. 4Pi-SMS images of Golgin-84 (magenta) and p230 (green). Arrows show the gap between Golgin-84 and p230 in side view.

E. Illustration of locations of Golgin-84, TMF, and Giantin relative to GALNT2 and p230.

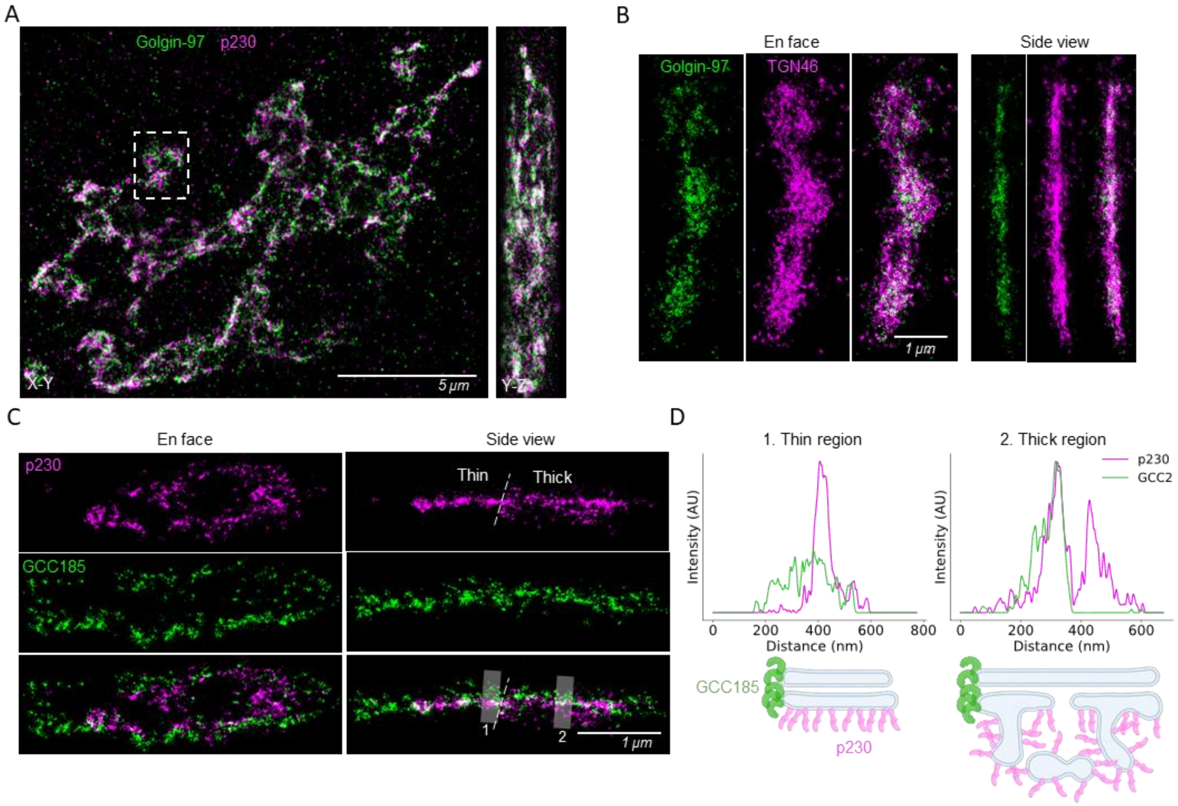


**Supplemental Figure 7. Relative localizations of *trans* golgins.**

A. 4Pi-SMS overview images of the entire Golgi region in a cell labeled by Golgin-97 and p230. White dashed box shows the region used in Fig. 4C.

B. Colocalization of TNG46 and Golgin-97 showed by 4Pi-SMS.

C. 4Pi-SMS images of sheet p230 and rim GCC185. Dashed line separates thin and thick regions in side view.

D. Top: Line-scan profiles of thin (Left) and thick (Right) regions. Bottom: illustrations of locations of GCC185 relative to p230.

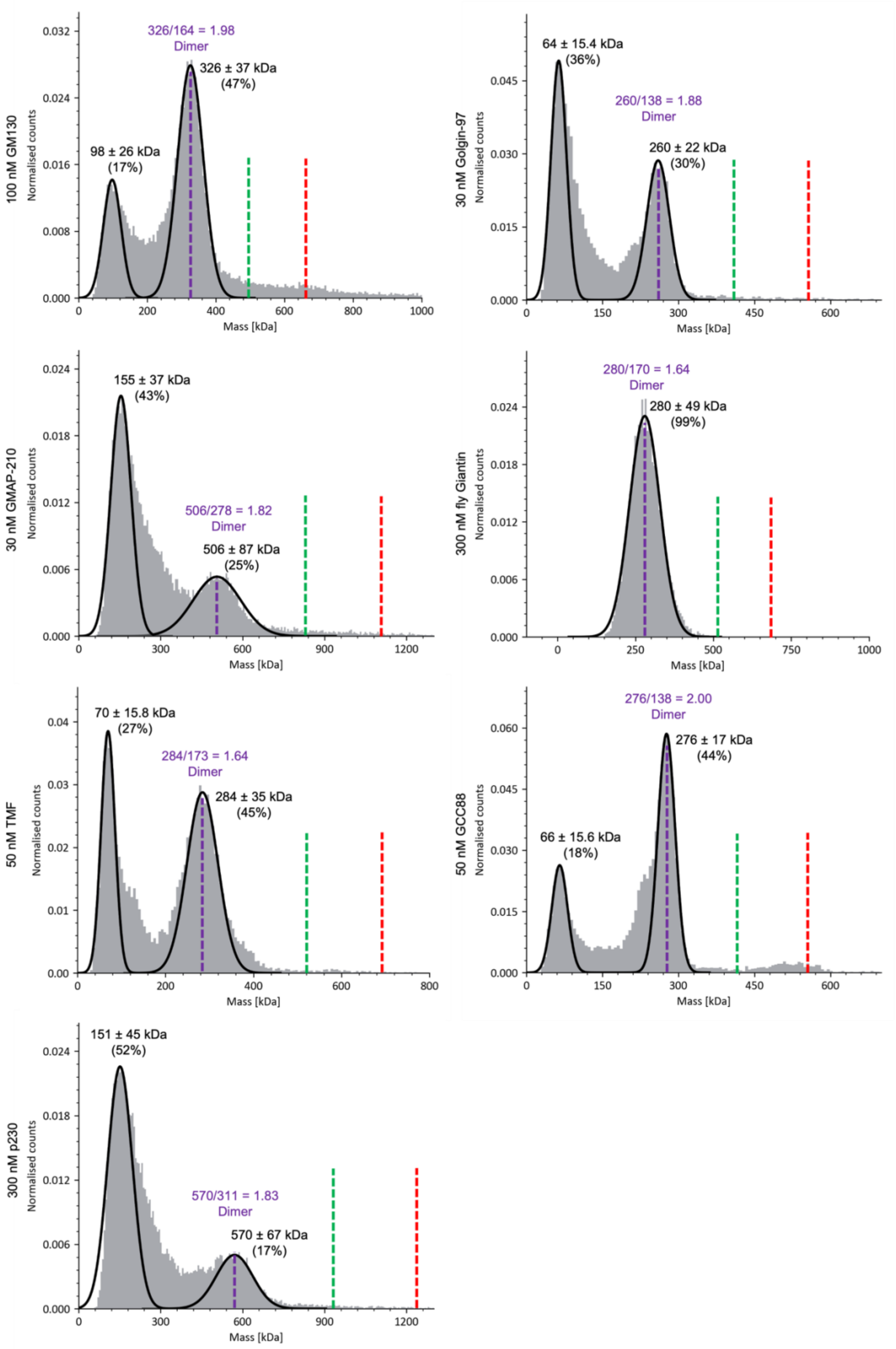
100 nM GM130
Normalised counts
326/164 = 1.98
Dimer
326 ± 37 kDa
(47%)
98 ± 26 kDa
(17%)
Mass [kDa]
30 nM Golgin-97
Normalised counts
64 ± 15.4 kDa
(36%)
260/138 = 1.88
Dimer
260 ± 22 kDa
(30%)
Mass [kDa]
30 nM GMAP-210
Normalised counts
155 ± 37 kDa
(43%)
506/278 = 1.82
Dimer
506 ± 87 kDa
(25%)
Mass [kDa]
300 nM fly Giantin
Normalised counts
280/170 = 1.64
Dimer
280 ± 49 kDa
(99%)
Mass [kDa]
50 nM TMF
Normalised counts
70 ± 15.8 kDa
(27%)
284/173 = 1.64
Dimer
284 ± 35 kDa
(45%)
Mass [kDa]
50 nM GCC88
Normalised counts
276/138 = 2.00
Dimer
276 ± 17 kDa
(44%)
66 ± 15.6 kDa
(18%)
Mass [kDa]
300 nM p230
Normalised counts
151 ± 45 kDa
(52%)
570/311 = 1.83
Dimer
570 ± 67 kDa
(17%)
Mass [kDa]

**Supplemental Figure 8. Mass photometry of purified golgins revealed the presence of dimers.**
Mass photometry measurements of purified full length golgin proteins with C-terminal bulky tags revealed dimers as the highest oligomeric species at low nanomolar concentrations. The highest oligomer peak is indicated by dashed purpled line. Purple label indicates the ratio between measured and theoretical molecular weight of the highest oligomeric peak. Dashed green and red lines indicate the estimated position of the trimer and tetramer peaks on the graph.

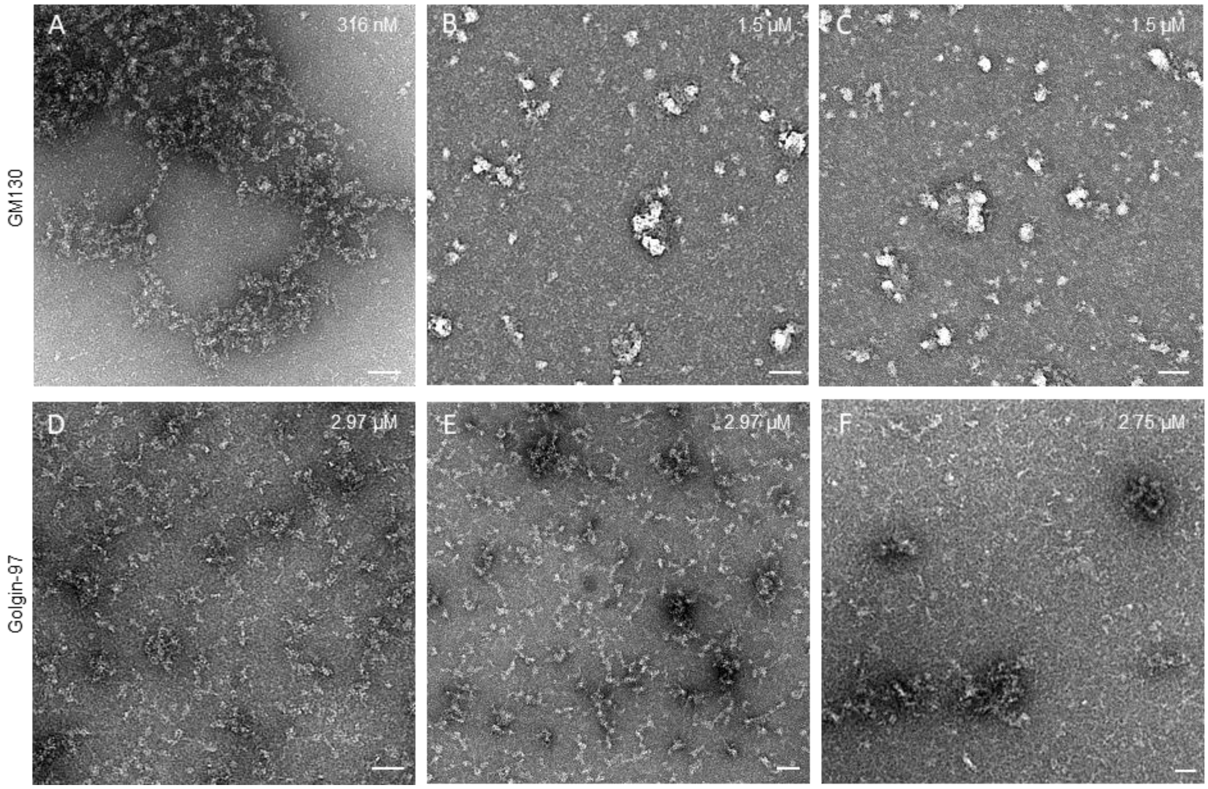


**Supplemental Figure 9. Negative stain EM images of irregular mesh like aggregates formed by sheet golgins.**
GM130 (A-C) and Golgin-97 (D-F) at high concentrations formed irregular mesh like aggregates on the carbon surface of the grid. Scale bar = 50 nm.

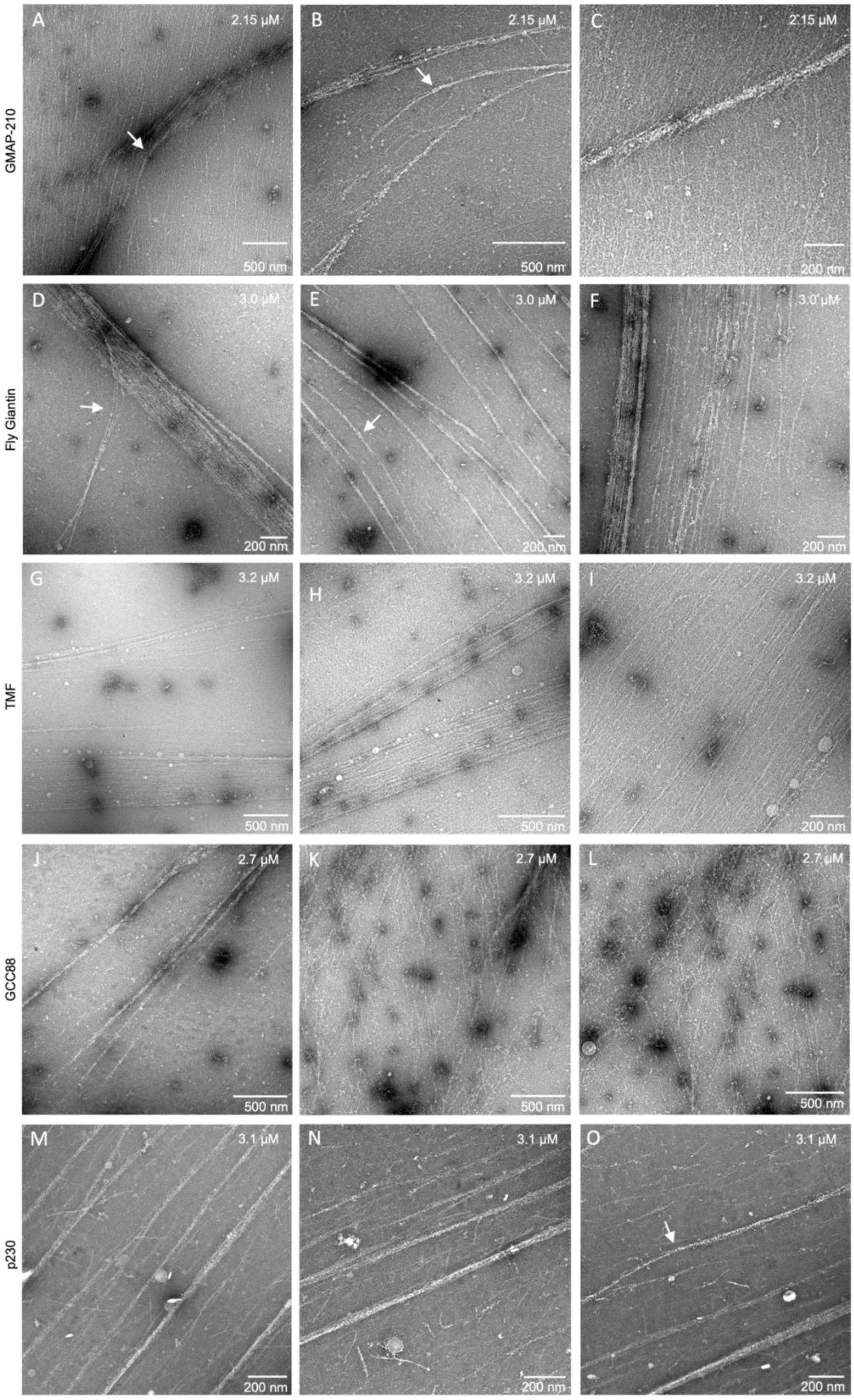
GMAP-210
Fly Giantin
TMF
GCC88
p230
A
2.15 μM
500 nm
B
2.15 μM
500 nm
C
2.15 μM
200 nm
D
3.0 μM
200 nm
E
3.0 μM
200 nm
F
3.0 μM
200 nm
G
3.2 μM
500 nm
H
3.2 μM
500 nm
I
3.2 μM
200 nm
J
2.7 μM
500 nm
K
2.7 μM
500 nm
L
2.7 μM
500 nm
M
3.1 μM
200 nm
N
3.1 μM
200 nm
O
3.1 μM
200 nm

**Supplemental Figure 10. Negative stain EM images of filament bundles formed by rim golgins.** GMAP-210 (A-C), melon fly Giantin (D-F), TMF (G-I), GCC88 (J-L) and p230 (M-O) show the presence of long filaments at micromolar concentrations on the carbon surface of the grid. White arrows point to the curved filament bands, wherever present.

### Supplemental Tables

| Name and epitope of primary antibody | Cat# | Company |
|---|---|---|
| rabbit Anti-GM130 antibody [EP892Y] (1-100 aa) | ab52649 | abcam |
| rabbit polyclonal anti-GM130 (685-1002 aa) antibody | 11308-1-AP | Proteintech |
| Mouse monoclonal anti-GM130 (869-982 aa) antibody IgG1, κ | 610823 | BD |
| Rabbit TRIP230 (GMAP-210 14-148 aa) Polyclonal Antibody | PA5-51726 | Invitrogen |
| GMAP-210 (1-68 aa) Rabbit Polyclonal antibody | 26456-1-AP | Proteintech |
| TRIP230 (GMAP-210 1099-1372 aa) Mouse Monoclonal IgG1 Antibody (10G5) | MA1-23294 | Invitrogen |
| TRIP11/TRIP230 (GMAP-210 1929-1979 aa) Rabbit Polyclonal Antibody | A301-188A | Bethyl |
| GOLGA5 Golgin-84 (159-279 aa) rabbit Polyclonal Antibody | HPA000992 | Sigma |
| GOLGA5 Golgin-84 (1-300 aa) rabbit Polyclonal Antibody | PA5-90689 | Invitrogen |
| CUX1/Protein CASP (1-341 aa) rabbit Polyclonal Antibody | 11733-1-AP | Proteintech |
| CUX1/Protein CASP rabbit Polyclonal Antibody (373-614 aa) | PA5-30003 | Invitrogen |
| CUX1/CASP (FL) purified MaxPab mouse polyclonal antibody (B01P) | H00001523-B01P | Abnova |
| TMF1 antibody produced in rabbit (253-397 aa) | HPA008729-100UL | Sigma |
| TMF1 mouse Monoclonal antibody (783-1093 aa) | 67505-1-Ig | Proteintech |
| TMF1 Recombinant Rabbit Monoclonal Antibody (JE65-73) (994-1,093 aa) | MA5-44831 | Invitrogen |
| Anti-GOLGB1 Giantin antibody rabbit (4-136 aa) | HPA011555-100UL | Sigma |
| Mouse anti-Giantin Antibody (2388C3a) (mapped to middle) | sc-81279 | Santa Cruz |
| Giantin (1250-1350 aa) Rabbit polyclonal antibody | ab80864 | Abcam |
| anti-Giantin (1462-1540 aa) antibody: scFv-TA10 mouse IgG2a | AA341-M2a | ABCD |
| Anti-GOLGB1 Giantin (1778-1906 aa) Rabbit Polyclonal Antibody | HPA011008 | Atlas Antibodies |
| Anti-Giantin Mouse monoclonal antibody [9B6] (mapped to C-term) | ab37266 | Abcam |
| Recombinant Anti-GRASP65 antibody [EPR12439] - C-terminal rabbit | ab174834 | Abcam |
| GRASP65 Antibody (OTI5G8) mouse | NBP2-02665 | Novus |
| GalNac Transferase 2/GALNT2 Antibody Polyclonal Sheep | AF7507-SP | Novus |
| Anti-GOLGA1 Golgin-97 (252-322 aa) Rabbit Polyclonal Antibody | HPA044329 | Atlas Antibodies |
| Golgin-97 rabbit Polyclonal antibody (418-767 aa) | 12640-1-AP | Proteintech |
| Anti-GOLGA1 Golgin-97 (659-765 aa) Rabbit Polyclonal Antibody | HPA050555 | Atlas Antibodies |
| Anti-Golgin-97 antibody Mouse polyclonal (1-800 aa) | ab169287 | Abcam |
| golgin-97 (GOLGA1 Full-length) mouse monoclonal IgG1 Antibody (CDF4) | sc-73619 | Santa Cruz |
| Rabbit polyclonal Anti Golgin-97 antibody (aa 750 to the C-terminus) | ab84340 | Abcam |
| GOLGA4 (p230) rabbit Polyclonal Antibody (a.a. 1-150) | PA5-87716 | Invitrogen |
| Rabbit Polyclonal Anti-GOLGA4 (p230) (1290-1380 aa) Antibody | HPA040675 | Atlas Antibodies |
| p230 (Golgin245, GOLGA4) (aa. 2063-2179) mouse antibody | 611281 | BD |
| GCC1 (264-343 aa) Rabbit Polyclonal antibody | PA5-53988 | Invitrogen |
| GCC1 (center region) Rabbit polyclonal Antibody | NBP2-16621 | Novus |
| GCC1 (GCC88 628-765 a.a.) antibody produced in rabbit | HPA021323-100UL | Sigma |
| GCC1 (full length) purified MaxPab mouse polyclonal antibody (B01P) | H00079571-B01P | Abnova |

| | | |
|---|---|---|
| anti-GCC2 (GCC185 1-300 a.a.) | PA5-89457 | Invitrogen |
| GCC2 (220-319 aa) Polyclonal Antibody rabbit | HPA035849 | Atlas Antibodies |
| Rabbit polyclonal GCC2 (1316-1558 aa) antibody | PAB15688 | Abnova |
| TGN46/TGOLN2 Rabbit mAb (1-100 aa) | A19618 | Abclonal |
| TGN46 (25-384 aa) mouse Monoclonal antibody | 66477-1-Ig | Proteintech |
| TGN46 rabbit Polyclonal antibody (a.a. 25-384) | 13573-1-AP | Proteintech |
| Rabbit anti Human TGN46 (full-length) Polyclonal Antibody serum | AHP1586 | Bio-rad |
| Sheep anti Human TGN46 (full-length) Polyclonal Antibody purified | AHP500G | Bio-rad |
| FLAG tag (DYKDDDDK) Mouse / IgG2b Monoclonal antibody | 66008-4-Ig | Proteintech |
| Rabbit polyclonal ant-GFP | A11122 | Invitrogen |
| Mouse Monoclonal GFP antibody | DSHB-GFP-4C9 | DSHB |

**Table S1. Primary antibodies used in this work.**

| Name and dye of secondary antibody | Cat# | Company |
|---|---|---|
| Goat anti-Mouse Alexa Fluor 647 | A21236 | Invitrogen |
| Goat anti-Rabbit Alexa Fluor 647 | A21245 | Invitrogen |
| Goat anti-Mouse CF660C for STORM | 20812-500µL | Biotium |
| Goat anti-Rabbit CF660C for STORM | 20813-500µL | Biotium |
| Goat anti-Rabbit IgG (H+L) Cross-Adsorbed Secondary Antibody, DyLight 633 | 35563 | Invitrogen |
| Goat anti-Mouse IgG (H+L) Cross-Adsorbed Secondary Antibody, DyLight 633 | 35513 | Invitrogen |
| Goat anti-Rabbit IgG (H+L) Cross-Adsorbed Secondary Antibody, DyLight 650 | #20818-500uL | Biotium |
| Goat anti-Mouse IgG (H+L) Cross-Adsorbed Secondary Antibody, DyLight 650 | #20817-500uL | Biotium |
| Donkey Anti-Sheep IgG H&L antibody DyLight 650 | ab96942 | Abcam |
| Goat Anti-Rabbit IgG (H+L), Highly Cross-Adsorbed Secondary Antibody, CF680 Single Label for STORM | #20818-500uL | Biotium |
| Goat Anti-Mouse IgG (H+L), Highly Cross-Adsorbed Secondary Antibody, CF680 Single Label for STORM | #20817-500uL | Biotium |
| Alexa Fluor® 647 AffiniPure Fab Fragment Goat Anti-Mouse IgG (H+L) | 115-607-003 | Jackson Immunoreseach |
| Alexa Fluor® 647 AffiniPure Fab Fragment Goat Anti-Rabbit IgG (H+L) | 111-607-003 | Jackson Immunoreseach |
| Donkey anti-Sheep IgG (H+L) Cross-Adsorbed Secondary Antibody, Alexa Fluor 647 | A-21448 | Invitrogen |
| Donkey anti-Goat IgG (H+L) Cross-Adsorbed Secondary Antibody, Alexa Fluor 647 | A-21447 | Invitrogen |
| FlexAble CoraLite® Plus 647 Antibody Labeling Kit for Rabbit IgG | KFA003 | Proteintech |
| Alexa Fluor® 647 VHH Fragment Alpaca Anti-Mouse IgG (H+L) | 615-604-214 | Jackson Immunoreseach |
| Alexa Fluor® 647 VHH Fragment Alpaca Anti-Rabbit IgG (H+L) | 611-604-215 | Jackson Immunoreseach |
| Alexa Fluor® 647 AffiniPure Fab Fragment Donkey Anti-Goat IgG (H+L) | 705-607-003 | Jackson Immunoreseach |

| | | |
|---|---|---|
| Donkey Anti-Rabbit IgG (H+L), Highly Cross-Adsorbed, CF-660C, Single Label for STORM | #20816-500uL | Biotium |
| Donkey Anti-Mouse IgG (H+L), Highly Cross-Adsorbed, CF-660C, Single Label for STORM | #20815-500uL | Biotium |
| Custom conjugation anti-mouse VHH (Jackson Immuno) with CF660C | custom | Biotium |
| Custom conjugation anti-rabbit VHH (Jackson Immuno) with CF660C | custom | Biotium |
| Donkey anti-Rabbit IgG (H&L), DyLight 633 conjugated, Cross-Adsorbed Secondary | AS12 2034 | Agrisera |
| Donkey anti-Mouse IgG (H&L), DyLight 633 conjugated, Cross-Adsorbed Secondary | AS12 2181 | Agrisera |
| Donkey anti-Rabbit IgG (H&L), DyLight 650 conjugated, Cross-Adsorbed Secondary | AS12 2329 | Agrisera |
| Donkey anti-Mouse IgG (H+L) DyLight 650 Cross-Adsorbed Secondary | SA5-10169 | Invitrogen |
| Donkey Anti-Sheep IgG (H+L) antibody, Highly Cross-Adsorbed CF680, single-label for STORM | Custom | Biotium |
| Abberior FLUX 680, donkey anti-sheep IgG, 500 µl (1 mg/ml) | FX680-1056-500UG | Abberior |

**Table S2. Secondary antibodies used in this work.**

| Layer | Golgin | FWHM[1] (nm) | Distance to GM130[2] (nm) |
|---|---|---|---|
| **1** | GM130 | 110±16 | N/A |
| **1** | GMAP-210 | 89±11 | 11±9 |
| **2** | Golgin-84 | 80±14 | 65±20 |
| **3** | TMF | 85±16 | 126±30 |
| **2,3** | Giantin | 159±37 | 93±27 |
| **2,3** | CASP | 134±32 | 78±28 |
| **4,5** | GCC88 | 137±31 | 228±48 |
| **4,5** | GCC185 | 168±33 | 202±36 |

**Table S3. Width and spacing of golgin layers. Corresponding to Fig. 2IJ.**

1. 5-11 cells, 13-33 ROIs for each protein. Shown as mean±SD.
2. 4-8 cells, 10-16 ROIs for each protein. Shown as mean±SD.

| Golgin | LQ[3] | Immuno-EM | Partition 4Pi-SMS | Cisterna No. 4Pi-SMS | Oligomer orientation at 10-150 nM Negative stain EM | Forming long filaments at μM Negative stain EM |
|---|---|---|---|---|---|---|
| GM130 | 0 | *cis*(Nakamura et al., 1995); *cis* side of *cis* sheet[1] | sheet | 1 | Parallel dimers | No |
| GMAP-210 | - | *cis* and vesicles (Pernet-Gallay et al., 2002) | rim | 1 | Parallel and anti-parallel dimers | Yes |
| Giantin | 0.57 | rim(Koreishi et al., 2013) | rim | 2, 3 | Parallel and anti-parallel dimers[2] | Yes[2] |
| CASP | - | | rim | 2, 3 | | |
| Golgin-84 | 0.26 | throughout(Satoh et al., 2003) | rim | 2 | | |
| TMF | - | rim(Yamane et al., 2007) | rim | 3 | Parallel and anti-parallel dimers | Yes |
| ManII | 0.23 | medial(Novikoff et al., 1983) | sheet | 2, 3, 4 | | |
| GALNT2 | 0.86 | medial(Röttger et al., 1998) | sheet | 2, 3 | | |
| GCC185 | 0.94 | *trans*(Luke et al., 2003) | rim | 4, 5 | Parallel dimers by AFM[4] | |
| GCC88 | - | *trans*(Luke et al., 2003) | rim | 4, 5 | Parallel and anti-parallel dimers | Yes |
| Golgin-245 /p230 | 1.42 | *trans*(Kooy et al., 1992) | sheet | 5+ | Parallel and anti-parallel dimers | Yes |
| Golgin-97 | 1.45 | | sheet | 5+ | Parallel dimers | No |

**Table S4. Summary and comparison of localization results for mutual Golgi proteins.**

Foot notes:

1. This work.
2. Data from melon fly giantin.
3. localization quotient (LQ) of averaged ministacks by Airyscan microscopy. Data from (Tie et al., 2018).
4. Atomic force microscopy (AFM) data from (Cheung et al., 2015).

### Supplemental Movies

Movie S1. An entire Golgi in HeLa cell labeled by GM130 (magenta) and Giantin (green).

Movie S2. Zoomed-in Golgi stacks labeled by GM130 (magenta) and GMAP-210 (green).

Movie S3. A zoomed-in Golgi stack labeled by GM130 (magenta)and GCC185 (green).

Movie S4. pan-ExM of GALNT2 and NHS shown in Z-slices.

Movie S5. 4Pi-SMS of p230_N (magenta) p230_C (yellow) showing thin and thick regions of *trans* Golgi.
